# A genetically informed atlas of transcriptomic programs from massive human RNA-seq compendia

**DOI:** 10.1101/2025.06.05.658122

**Authors:** Marc Subirana-Granés, Haoyu Zhang, Sutanu Nandi, Jill Hoffman, Sakaiza Rasolofomanana-Rajery, Rawan Elshobaky, Lucas A. Gillenwater, Alexander P. McKim, James C. Costello, Maria Chikina, Milton Pividori

## Abstract

Genetic risk for complex disease acts through coordinated gene programs in specific tissues and cellular contexts, which single-gene analyses cannot resolve. Interpretable gene modules can capture these programs, but prior-informed methods have not scaled to large expression compendia, and genetic associations have not been integrated with modules learned at this scale. Here we present CLAMP, a fast, memory-efficient prior-informed framework that learns interpretable gene modules from such compendia and integrates them with genetic associations and drug signatures. CLAMP recovered cell-type composition and tissue organization more accurately than existing decomposition methods. Training on 605,614 ARCHS4 samples increased pathway and genetic-trait recovery, and projection into these modules recovered programs missed by study-specific models. ARCHS4 modules also ranked known drug-disease pairs better than modules learned from smaller compendia or individual genes. In reproducible consensus modules, cell-type, pathway, drug-response and genetic evidence converged on shared processes, placing CRISPR-validated lipid regulators missed by single-gene tests in disease-associated programs. CLAMP is available through Bioconductor, and its trained models provide a genetically informed atlas of gene programs for interpreting complex disease.

## Introduction

Modern models of complex disease propose that genetic risk acts through regulatory networks and coordinated molecular programs^1–3^. Because genetic regulatory effects depend on tissue, cell type and cell state, interpreting their contribution to disease requires understanding how they alter molecular programs and cellular functions in these contexts^4–6^. Transcriptome-wide association studies (TWAS) test associations between genetically predicted expression and traits to prioritize candidate genes^7,8^. However, single-gene approaches do not explicitly model gene-gene interactions, limiting the identification of coordinated programs implicated in diseases^9^. Co-expression modules learned across diverse tissues and cell populations could connect disease-associated genes to coordinated transcriptional programs that reflect shared regulation and function.

Unsupervised decomposition methods provide a framework for learning such co-expression modules and have outperformed other module-detection methods^10^. Within this framework, some approaches incorporate biological priors to improve interpretability. PLIER pulls gene loadings toward combinations of curated pathways^11^, and MOFA-FLEX assigns predefined gene sets to individual factors^12^. Because both use gene sets to shape module composition directly, they can underweight co-expressed genes outside existing annotations, and their computational demands have limited their application to compendia far smaller than current public resources. Transfer learning extends the utility of these methods by applying programs learned from large expression collections to independent datasets. MultiPLIER demonstrated that modules learned from a heterogeneous compendium can reveal biological signals in smaller studies and improve understanding of processes associated with disease^13^. GenomicSuperSignature constructed replicable axes of variation from principal components across public studies, supporting interpretation of new RNA-seq datasets^14^.

PhenoPLIER further demonstrated that pretrained expression modules can connect genetic associations with drug-induced gene expression and prioritize functionally supported genes^15^. Together, these studies show that interpretable expression modules can capture biological signals across independent datasets and connect genetic associations with perturbational responses to improve our understanding of disease biology.

Public resources such as ARCHS4 now provide large compendia of uniformly processed human RNA-seq profiles spanning tissues, cell types, diseases and treatments^16^. Their diversity offers opportunities to recover cell-and tissue-specific programs, as well as treatment responses and other biological processes poorly captured by models trained on smaller expression compendia. However, it remains unclear whether training prior-informed models on larger compendia produces modules that capture a broader range of biological processes, retain cell-and tissue-specific signals, and better support transfer learning and interpretation of genetic associations.

Here we present CLAMP (Curated Latent-variable Analysis with Molecular Priors), a scalable framework that learns interpretable transcriptional programs from large expression compendia, integrates them with genetic and perturbational data and transfers them across datasets, providing a way to operationalize modern models of complex disease. CLAMPbase learns gene modules using semi-nonnegative matrix factorization, whereas CLAMPfull uses curated gene sets as molecular priors to adjust how strongly each gene’s loading is shrunk, guiding modules toward known pathways while retaining genes supported only by the expression data. Algorithmic optimizations and file-backed computation enable training on more than 600,000 ARCHS4 samples, and new expression datasets can be projected into the learned module space, where they can be analysed together with genetic associations and drug perturbations. We show that CLAMP resolves cell-type and tissue programs more accurately than existing decomposition methods, including in bulk data, and that training on the full compendium broadens the pathways, traits and study-specific programs its modules capture.

Applied to genetic and drug-perturbation data, these modules improve drug-disease prediction and connect experimentally validated genes without individual TWAS signals to disease. We release CLAMP as a Bioconductor package and the trained models as a browsable, genetically informed atlas of transcriptional programs.

## Results

### Overview of CLAMP

CLAMP is a matrix factorization framework that learns interpretable gene modules (GMs) from large expression compendia and connects them to pathways, cell types, drug-induced transcriptional responses and genetic associations (Figure 1). Given a gene-by-sample expression matrix *Y* , CLAMP approximates it as the product *ZB*, where the gene loadings *Z* (genes × GMs) quantify each gene’s contribution to each GM and the module activities *B* (GMs × samples) quantify each GM’s activity across samples (Figure 1a).

**Figure 1:**
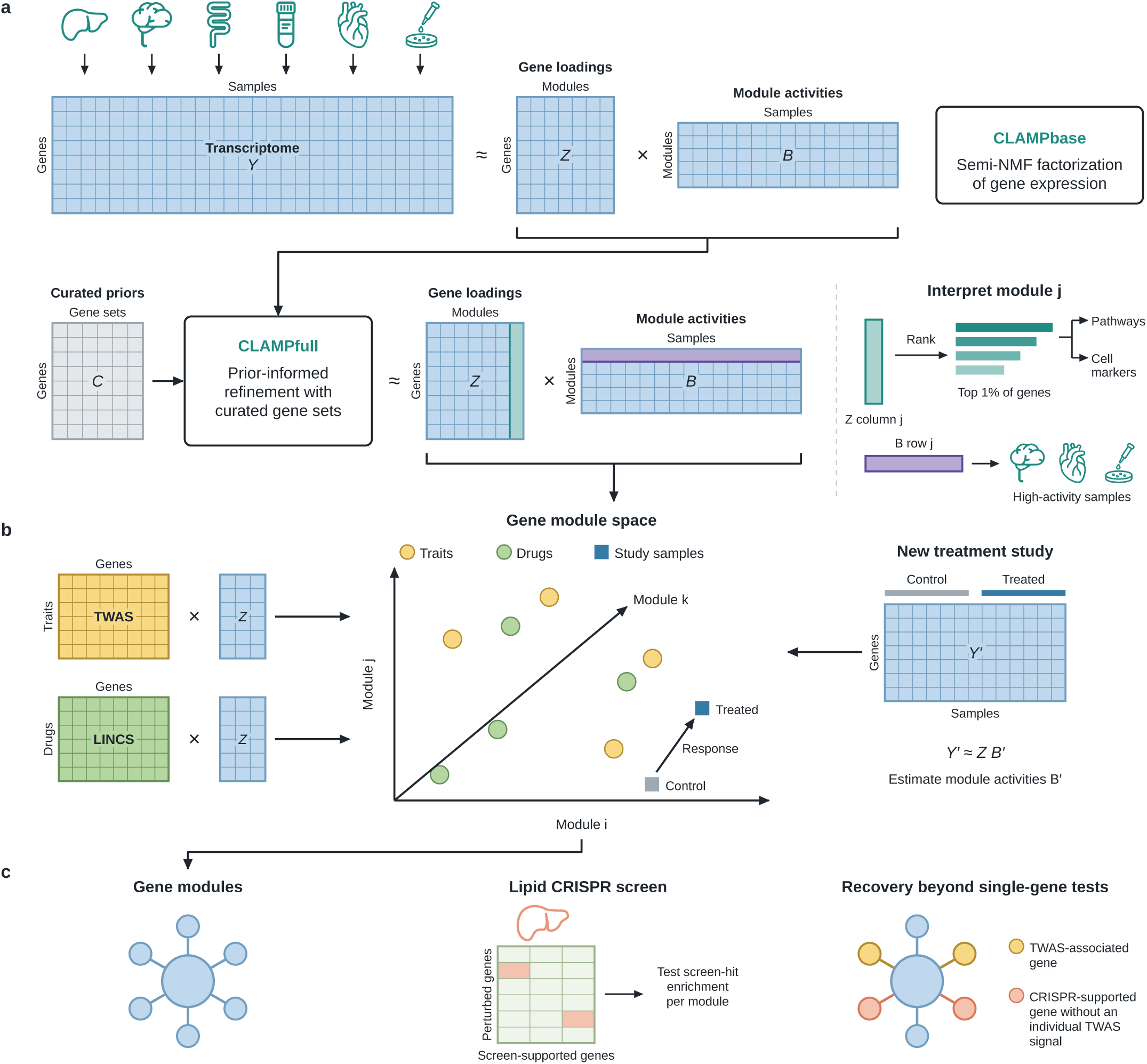
CLAMP learns interpretable gene modules as a common reference for expression data, drug-induced transcriptional responses and genetic associations. a, Model learning and interpretation. A gene-by-sample expression matrix *Y* is approximated by the product of gene loadings *Z* (genes by gene modules (GMs)) and module activities *B* (GMs by samples). CLAMPbase learns GMs using semi-nonnegative matrix factorization (semi-NMF). CLAMPfull is initialized from this solution and uses curated gene sets, represented by the gene-by-gene-set matrix *C*, to identify the gene sets associated with each GM and guide a variance prior on gene loadings. For GM *j*, the top 1% of genes ranked by loading in column *j* of *Z* are used for pathway and cell-marker enrichment analyses. High-activity samples in row *j* of *B* identify tissue, cellular and experimental contexts. *b*, Transfer learning and projection into a shared gene module space. New expression data *Y ′* are projected into the learned gene module space without retraining, estimating module activities *B′* with the gene loadings *Z (Y ′ ≈ ZB′)*. Gene-level transcriptome-wide association study (TWAS) and Library of Integrated Network-based Cellular Signatures (LINCS) drug perturbation signatures are mapped into the same space using the learned gene loadings, adapting the PhenoPLIER framework^15^. Yellow circles, traits; green circles, drugs; squares, study samples. c, Evaluation using a published lipid CRISPR screen. Enrichment of screen-supported genes in trait-associated GMs assesses recovery of genes with experimental evidence for lipid regulation, including genes without individual TWAS signals for the corresponding traits. Yellow nodes, TWAS-associated genes; salmon nodes, CRISPR-supported genes without individual TWAS signals. Matrices, networks and gene-module-space coordinates are schematic.

CLAMP is trained in two phases. CLAMPbase first estimates *Z* and *B* from expression data using semi-nonnegative matrix factorization. CLAMPfull is then initialized from the CLAMPbase solution and incorporates curated gene sets, represented as a gene-by-gene-set membership matrix *C*, as prior knowledge. For each GM, CLAMPfull predicts its gene loadings from *C* using cross-validated sparse regression, which selects the gene sets associated with that GM. These predictions set an adaptive variance prior on the gene loadings: genes supported by the selected gene sets are shrunk less toward zero, whereas all other genes are shrunk more strongly. This regularization guides GMs toward known biological programs while retaining patterns supported by the expression data beyond the predefined gene sets.

Once trained, each GM can be interpreted through its highest-loading genes and the samples in which it is most active (Figure 1a). New expression datasets can also be projected into the learned gene module space by estimating their module activities from the learned gene loadings, without retraining the model (Figure 1b), so that GMs learned from large reference compendia can be applied to smaller datasets^13^. Using the PhenoPLIER framework^15^, CLAMP integrates gene-level TWAS associations and drug-induced transcriptional profiles into the same GM space, enabling a joint analysis of genetic traits, drugs and expression samples (Figure 1b). GMs supported by several data types can then be prioritized for experimental validation, for example with perturbation screens (Figure 1c).

CLAMP trains faster and with lower memory requirements than other prior-informed approaches (Supplementary Figure 4) by combining algorithmic and computational optimizations. Gene sets are introduced only after an unsupervised phase has captured the main structure of the data, the regularization for each module is tuned efficiently, and file-backed expression matrices are processed blockwise so that training is not limited by memory. This scalability allowed us to build a reference atlas of GMs from virtually all public human bulk RNA-seq data in ARCHS4 (605,614 samples), in which each GM is annotated with pathways, cell types, drug perturbations and genetic traits.

## CLAMP recovers cell-type composition and cell-type-associated programs across pseudobulk and bulk RNA-seq

We first asked whether CLAMP could recover cell-type-associated signals when cell-type composition was known. To test this, we generated donor-level pseudobulk profiles from six single-cell or single-nucleus RNA-seq datasets spanning peripheral blood^17,18^, brain^19,20^, lung^21^ and heart^22^ by averaging cell-level expression within each donor while retaining cell-type proportions from the original annotations. We benchmarked CLAMP against state-of-the-art matrix decomposition methods, including PLIER^11^, MOFA-FLEX^12^, flashier^23^, GenomicSuperSignature (GSS)^14^ and CoGAPS^24^, with PCA^25^, NMF^26^ and ICA^27^ as baselines. To avoid circularity, prior-informed methods (CLAMPfull, PLIER and MOFA-FLEX) were trained on Gene Ontology Biological Process (GO:BP), which overlapped minimally with the marker sets used for evaluation (Supplementary Figure 1c). Modules were assigned to cell types by greedy one-to-one matching on Pearson correlation with the known proportions (Methods).

Across 47 dataset-cell-type targets, CLAMPfull and CLAMPbase achieved the highest mean correlations (r = 0.765 and 0.755; Figure 2a). Their mean performance did not differ significantly ( q = 0.383), but CLAMPfull ranked first among all ten methods in 94.9% of 2,000 dataset-level bootstrap resamples. Either CLAMP variant also achieved the highest mean correlation in each of the four tissues (Supplementary Figure 1a,b). To determine whether this performance generalized to unseen donors, we used fivefold donor-grouped cross-validation, assigning each cell type a module using training donors only. Held-out expression was projected onto the training-fold gene loadings, and a univariate linear regression fitted to training donors calibrated the assigned module’s activity to a proportion (Methods). For the 35 of 47 targets with a training correlation of at least 0.5 in every fold, mean prediction accuracy was r = 0.823, with a mean absolute error (MAE) of 0.040 (Figure 2b). Thus, CLAMP recovered cell-type composition and retained predictive accuracy in held-out donors.

**Figure 2:**
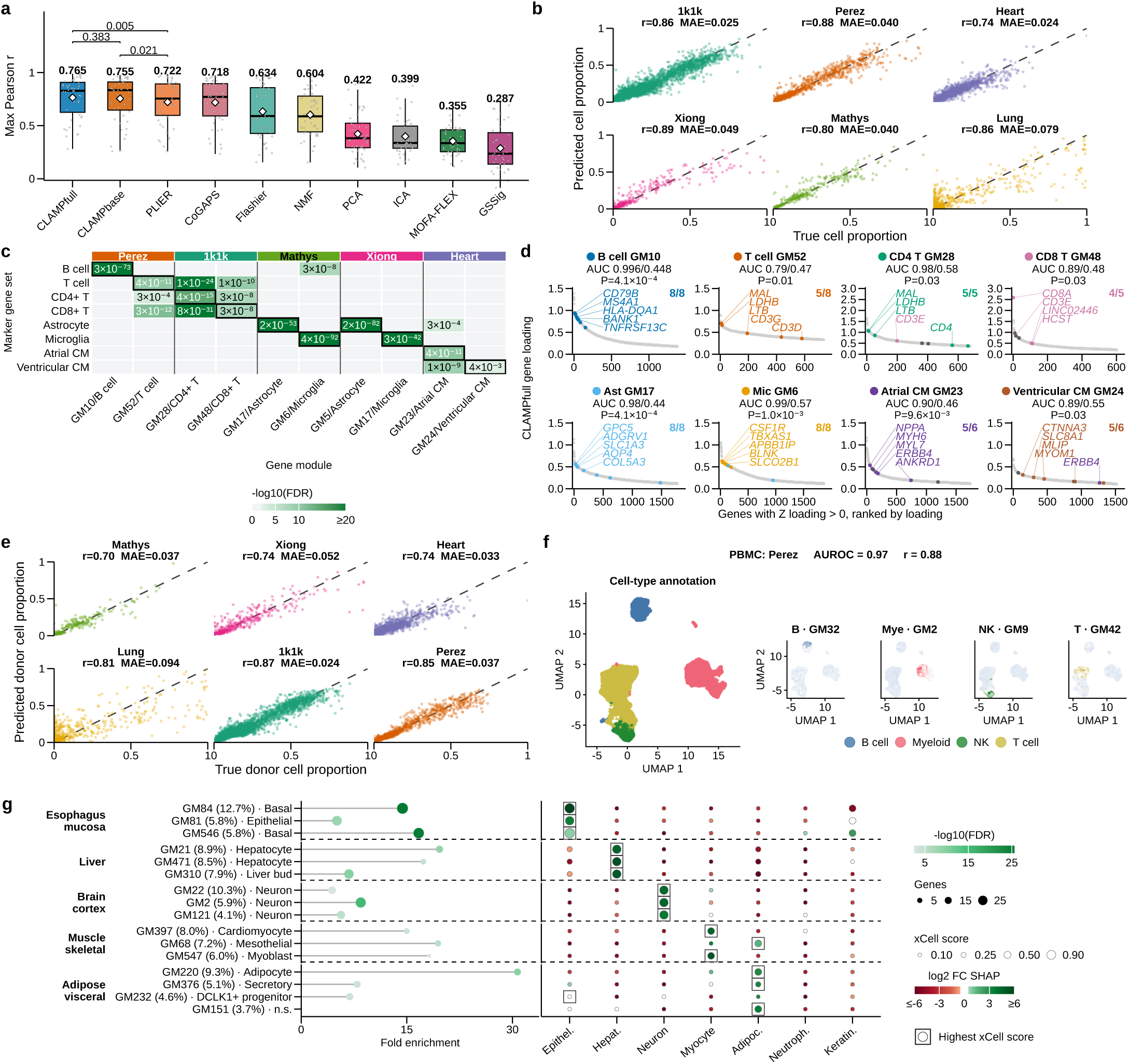
CLAMP recovers cell-type composition and cell-type-associated programs across pseudobulk and bulk RNA-seq. a, Cell-type proportion recovery for 47 dataset-cell-type targets, measured as the maximum Pearson correlation between known proportions and module activities after greedy one-to-one matching (Methods). Points, targets; boxes, median and interquartile range; whiskers, 1.5× interquartile range; diamonds and labels, means; brackets, Benjamini-Hochberg-adjusted two-sided paired Wilcoxon tests. b, Known versus predicted proportions for 35 stable targets in donor-grouped fivefold cross-validation; points, pooled out-of-fold predictions; labels, cohort mean Pearson *r* and mean absolute error (MAE); dashed lines, identity. c, Over-representation analysis (ORA) of the top 1% of genes by signed loading for 10 modules of eight closely related cell types, tested against all eight marker sets. Colour, − log10(FDR), capped at 20, with FDR corrected within each marker collection; gray, FDR ≥ 0.01; values, rounded FDR; outlines, matched module-marker pairs. d, Loading ranks of genes with positive *Z* loadings for the cell types in c; coloured points, markers exclusive to each member of a pair; fractions, exclusive markers with positive loading; titles, assigned-versus partner-marker AUROC and two-sided Wilcoxon rank-sum *P* . e, Known versus predicted proportions from donor-level count-summed bulk-like profiles for 32 stable targets; cross-validation, points and labels as in b. f, Perez PBMC cells projected into modules learned from donor-level count-summed profiles: cell-type annotations on an independent expression UMAP (left) and module activities (right); labels, median cell-level AUROC and donor-level Pearson *r* across four cell types; other datasets, Supplementary Figure 2b-f. g, Cell-type associations of 16 SHAP-prioritized GTEx modules across five subtissue-cell-type pairs; rows, subtissue, module, SHAP contribution and expected marker set. Left, enrichment of the top 1% of loadings for the expected CellMarker 2.0 markers; position, fold enrichment; colour, − log_10_(FDR); size, overlap. Right, mean xCell scores of seven cell types in each module’s top 1% highest-activity samples; size, mean score; outlines, highest score per module; fill, log2 tissue SHAP fold change. AUROC, area under the receiver operating characteristic curve; SHAP, Shapley additive explanations; UMAP, uniform manifold approximation and projection; GM, gene module; CM, cardiomyocyte; NK, natural killer; Mye, myeloid; Epithel., epithelial; Hepat., hepatocyte; Adipoc., adipocyte; Neutroph., neutrophil; Keratin., keratinocyte.

Beyond recovering composition, we asked whether CLAMP models trained on the full datasets captured the expected cell-type transcriptional programs. Gene loadings (Z) were evaluated against independent marker sets from Azimuth^28^, CellMarker 2.0^29^ and, for brain datasets, the Allen Brain Atlas^30^. Over-representation analysis (ORA) recovered the matched cell type for 39 of 47 modules (83.0%; Supplementary Figure 1e). Among eight closely related cell types, B cells, T cells, CD4+ T cells, CD8+ T cells, astrocytes, microglia, and atrial and ventricular cardiomyocytes, the corresponding modules tracked the expected cell-type proportions (r = 0.78 to 0.96; Figure 2c). Every matched comparison across the five datasets was significantly enriched for the expected marker set (FDR < 0.01; Figure 2c). At the gene level, markers of the assigned cell type consistently ranked above those of its paired cell type across all eight cell types (AUROC 0.79 to 0.996 versus 0.44 to 0.58; Wilcoxon rank-sum P = 0.0004 to 0.031; Figure 2d), including *CD79B* and *MS4A1* for B cells, *CD4* and *MAL* for CD4+ T cells, *CD8A* for CD8+ T cells, *GPC5* for astrocytes, *CSF1R* for microglia, *NPPA* and *MYL7* for atrial cardiomyocytes, and *CTNNA3* and *MYOM1* for ventricular cardiomyocytes. To test whether these programs were active in the expected cells, we projected the original single cells into the learned module space using a ridge-like regression that uses the gene loadings (Z) to estimate module activities (B) in new data. Cells with the matched annotation represented 76.8% of the top 1% highest-activity cells on average, corresponding to a mean 19.1-fold enrichment over expectation (Supplementary Figure 1d). Together, these analyses show that modules associated with cell-type proportions also encode the corresponding cell-type transcriptional programs in their gene loadings and sample activities.

Because conventional bulk RNA-seq mixes expression across cell types, we asked whether these signals remained detectable after aggregation. We therefore summed raw counts over all cells per donor regardless of cell type to generate overall pseudobulk profiles as a proxy for bulk tissue RNA-seq, while retaining the known underlying cell-type composition. Repeating module assignment, projection and calibration in donor-grouped fivefold cross-validation yielded mean r = 0.792 and MAE = 0.046 across 32 of 47 stable targets (Figure 2e). Projection of the original single cells onto the donor-bulk gene loadings recovered the corresponding cell types (median AUROC = 0.99; 44/47 ≥ 0.7; assigned cell type top-ranked for 39/47 modules; Figure 2f; Supplementary Figure 2a-f). Thus, cell-type-associated programs remained recoverable after aggregation into bulk-like expression profiles.

We then extended the analysis to GTEx bulk RNA-seq^4^, where the underlying cellular composition is unknown. CLAMPfull achieved the highest median adjusted Rand index (ARI) against GTEx subtissue labels among the benchmarked state-of-the-art methods (ARI = 0.634; Supplementary Figure 3b), exceeding the expression baseline, which performed best when restricted to the 25% most variable genes (Supplementary Figure 3a). We ranked modules for each subtissue using one-versus-rest random forest classifiers^31^ and Shapley additive explanations (SHAP)^32^. The gene loadings of the top-ranked module were enriched for the expected GTEx tissue gene set in 44 of 49 subtissues (Supplementary Figure 3c), and the module representation distinguished subtissues in held-out donors with balanced accuracies of 0.860 to 0.996 across seven tissues (Methods). These results show that CLAMP captures tissue and subtissue structure in conventional bulk RNA-seq.

To determine whether GTEx modules also captured cell-type-associated programs, we used xCell to provide sample-level estimates of cell-type enrichment in bulk RNA-seq and focused on five subtissue-cell-type pairs with available xCell scores: epithelial cells in esophagus mucosa, hepatocytes in liver, neurons in brain cortex, myocytes in skeletal muscle and adipocytes in visceral adipose tissue^33,34^.

For these pairs, we used SHAP values from one-versus-rest random forest classifiers to identify 16 modules that contributed most strongly to distinguishing the corresponding subtissues. Of these, 15 of 16 modules were enriched for markers of the expected cell type. Among samples in the top 1% of activity for each module, the expected cell type had the highest mean xCell score of the seven cell types tested for 14 of 16 modules (Figure 2g). Although xCell scores are themselves derived from bulk expression and do not provide independent single-cell validation, their agreement with marker enrichment supports the inferred cell-type associations.

Together, these results show that CLAMP recovers cell-type-associated programs from heterogeneous expression profiles across single-cell-derived pseudobulk, bulk-like overall pseudobulk and conventional bulk RNA-seq.

## Increasing training data scale broadens pathway and genetic-trait recovery

We next asked whether expanding the training compendium improves pathway and genetic-trait recovery. Previous work showed that models trained on larger and more heterogeneous expression compendia capture more biological programs than models trained on individual datasets, but these analyses were limited to compendia of tens of thousands of samples^13^. The scalability of CLAMP allowed us to extend this question to virtually all publicly available human bulk RNA-seq samples in ARCHS4^16^, and to ask how pathway and genetic-trait recovery change with training size. We progressively added complete studies in three subsampling series spanning 1% to 75% of ARCHS4 and compared these models with those trained on the complete ARCHS4 and GTEx compendia. We used unsupervised CLAMPbase without prior information to avoid circularity in recovering pathways and genetic-trait associations (Methods). For each training set, we set the number of modules (*K*) to twice the number of components estimated with the Gavish-Donoho method^35^ (Methods).

We first asked whether increasing the size of the training compendium improved the recovery of biological pathways. We tested enrichment of MSigDB canonical pathways^36^ and CellMarker cell-type gene sets^29^ among the top 1% of gene loadings of each module using over-representation analysis (ORA; FDR < 0.05; Methods). Median recovery increased from 1,364 of 2,010 eligible gene sets (68%) in models trained on 1% of ARCHS4 (6,109 samples) to 1,793 (89%) in models trained on the complete compendium (605,614 samples; Figure 3a). The complete ARCHS4 models recovered significantly more gene sets than models trained on 75%, 50% and 25% of the studies, with average gains of 37, 53 and 97 gene sets, respectively (P = 0.003, P = 0.004 and P = 6.8 × 10−4). They also recovered significantly more gene sets than GTEx-trained models (P = 0.007), with an average gain of 89 and a higher median recovery (1,793 versus 1,717; 89% versus 85%; Figure 3a).

**Figure 3:**
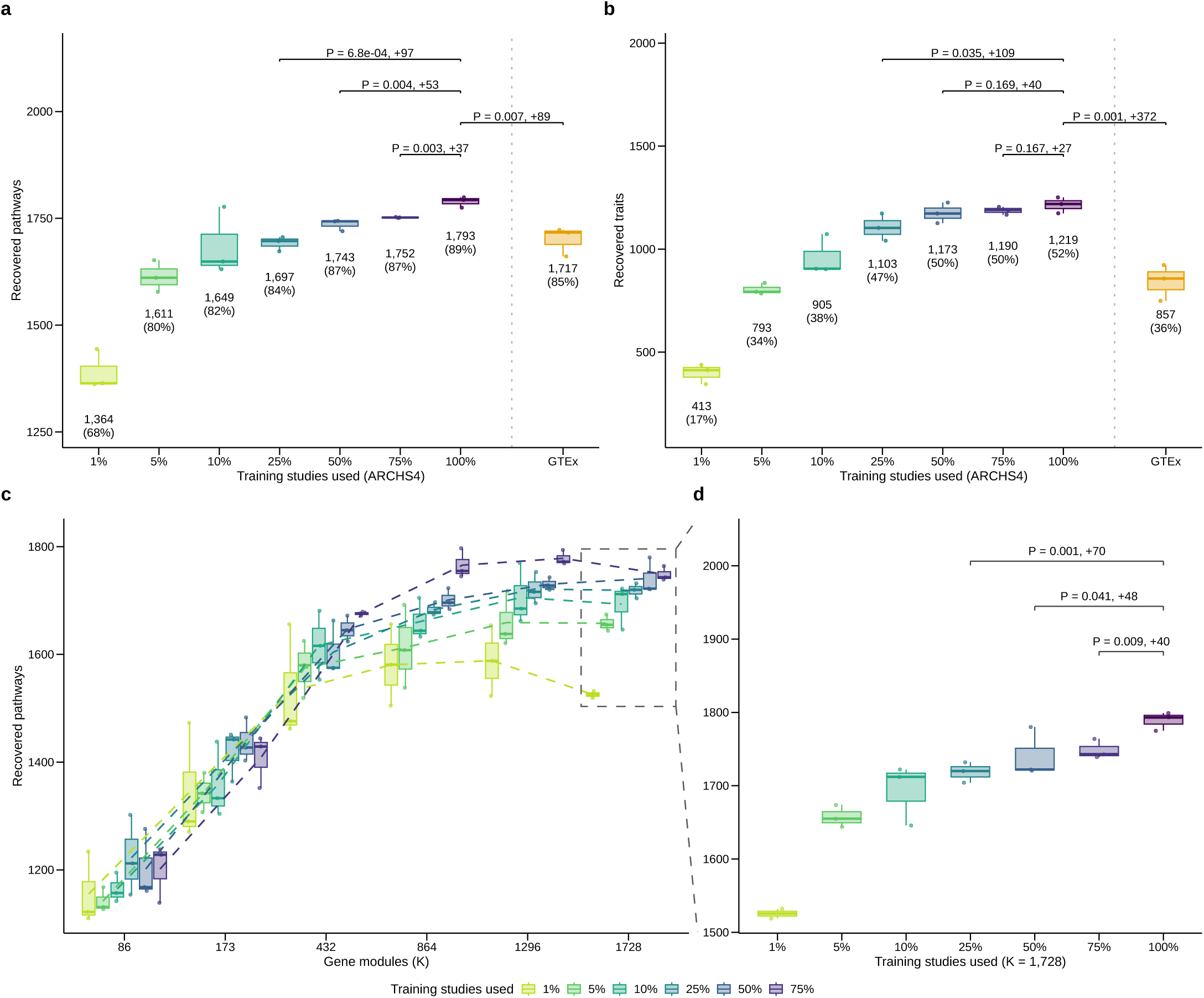
Larger training compendia recover more pathways and genetic-trait associations. a, Pathways recovered by unsupervised CLAMPbase models trained on study-level subsamples of ARCHS4 (left; *n* = 3 subsamples per fraction) and on the complete ARCHS4 and GTEx compendia (right; three independently fitted models each), labelled as median counts and as a percentage of the eligible gene sets (2,010 for ARCHS4 models and 2,024 for GTEx). Points, independent models; boxes, medians and interquartile ranges; whiskers, observations within 1.5 interquartile ranges. Brackets, the mean gain of the 100% ARCHS4 models over the labelled 75%, 50%, and 25% fractions and GTEx, with one-sided Student’s *t*-test *P* values. b, Traits recovered at false discovery rate (FDR) <0.05 by the same models, labelled as median counts and as a percentage of the 2,366 tested traits (Supplementary Table 1). Points, independent models; boxes, medians and interquartile ranges; whiskers, observations within 1.5 interquartile ranges. Brackets show the same 100% ARCHS4 comparisons as in a. c, Pathways recovered by CLAMPbase models retrained on each ARCHS4 subsample at six fixed numbers of gene modules (*K*; *n* = 3 subsamples per fraction and *K*, except *n* = 2 at 1% and *K* = 1,728; Methods); dashed box, the *K* = 1,728 group enlarged in d. Colours identify training fractions; points, subsamples; boxes, medians and interquartile ranges; whiskers, observations within 1.5 interquartile ranges; dashed lines, the mean of each fraction across *K*. d, The *K* = 1,728 models from c together with the three complete ARCHS4 models from a, whose estimated number of modules is 1,728, holding module number constant across training fractions (*n* = 3 per fraction, except *n* = 2 at 1%). Points, independent models; boxes, medians and interquartile ranges; whiskers, observations within 1.5 interquartile ranges. Brackets, the mean gain of the 100% models over the labelled 75%, 50% and 25% fractions and one-sided Student’s *t*-test *P* values.

We next examined whether these gains extended to genetic-trait associations using PhenoPLIER^15^, which applies generalized least-squares regression to test whether the top 1% of genes in each module show stronger transcriptome-wide association study (TWAS) associations with a trait than other genes. We evaluated 2,366 UK Biobank traits from PhenomeXcan^37^ and considered a trait recovered when at least one module was significantly associated with it (FDR < 0.05; Supplementary Table 1). Median trait recovery increased from 413 (17%) in models trained on 1% of ARCHS4 to 1,219 (52%) in models trained on the complete compendium (Figure 3b). The complete models recovered significantly more traits than the 25% models (109 additional traits on average; P = 0.035), and also recovered more traits than the 50% and 75% models, although these differences were not statistically significant (40 and 27 additional traits; P = 0.169 and P = 0.167, respectively). Compared with GTEx, the complete ARCHS4 models recovered significantly more traits (P = 0.001), with an average gain of 372 and a 42% increase in median recovery (1,219 versus 857; 52% versus 36%; Figure 3b).

Because the estimated number of modules increased with training-set size, we assessed whether the gains in recovery reflected training size or module number by retraining CLAMPbase on each subsample at six fixed values of K spanning 86 to 1,728. Although the estimated number of modules increased from K = 134 at 1% of ARCHS4 to K = 1, 728 in the complete compendium, pathway recovery continued to increase with training size even when K was held constant (Figure 3c). At K = 1, 728, the rank estimated for the complete compendium, the complete ARCHS4 models recovered a median of 1,793 pathways, significantly more than models trained on 75%, 50% and 25% (an average of 40, 48 and 70 more; P = 0.009, P = 0.041 and P = 0.001; Figure 3d). Trait recovery at fixed K = 1, 728 followed a similar trend, increasing from a median of 914 traits (39%) at 1% of ARCHS4 to 1,196 (51%) at 75% and 1,219 (52%) with the complete compendium. The complete models also recovered more traits than models trained on 75%, 50% and 25% (an average of 19, 49 and 42 additional traits, respectively), although these differences were not statistically significant (P = 0.258, P = 0.095 and P = 0.086; Supplementary Figure 5).

Together, these results show that larger training compendia improve the recovery of biological pathways and genetic-trait associations, with pathway gains persisting independently of the number of modules.

## Projection into the ARCHS4 model reveals biological programs missed by models trained on individual studies

We next asked whether a model trained on a large, heterogeneous compendium such as ARCHS4 could recover biological programs not captured by smaller or more homogeneous individual studies. CLAMP supports this through transfer learning: new expression data are projected into the ARCHS4 modules to estimate module activities for the new samples without retraining (Figure 4a).

**Figure 4:**
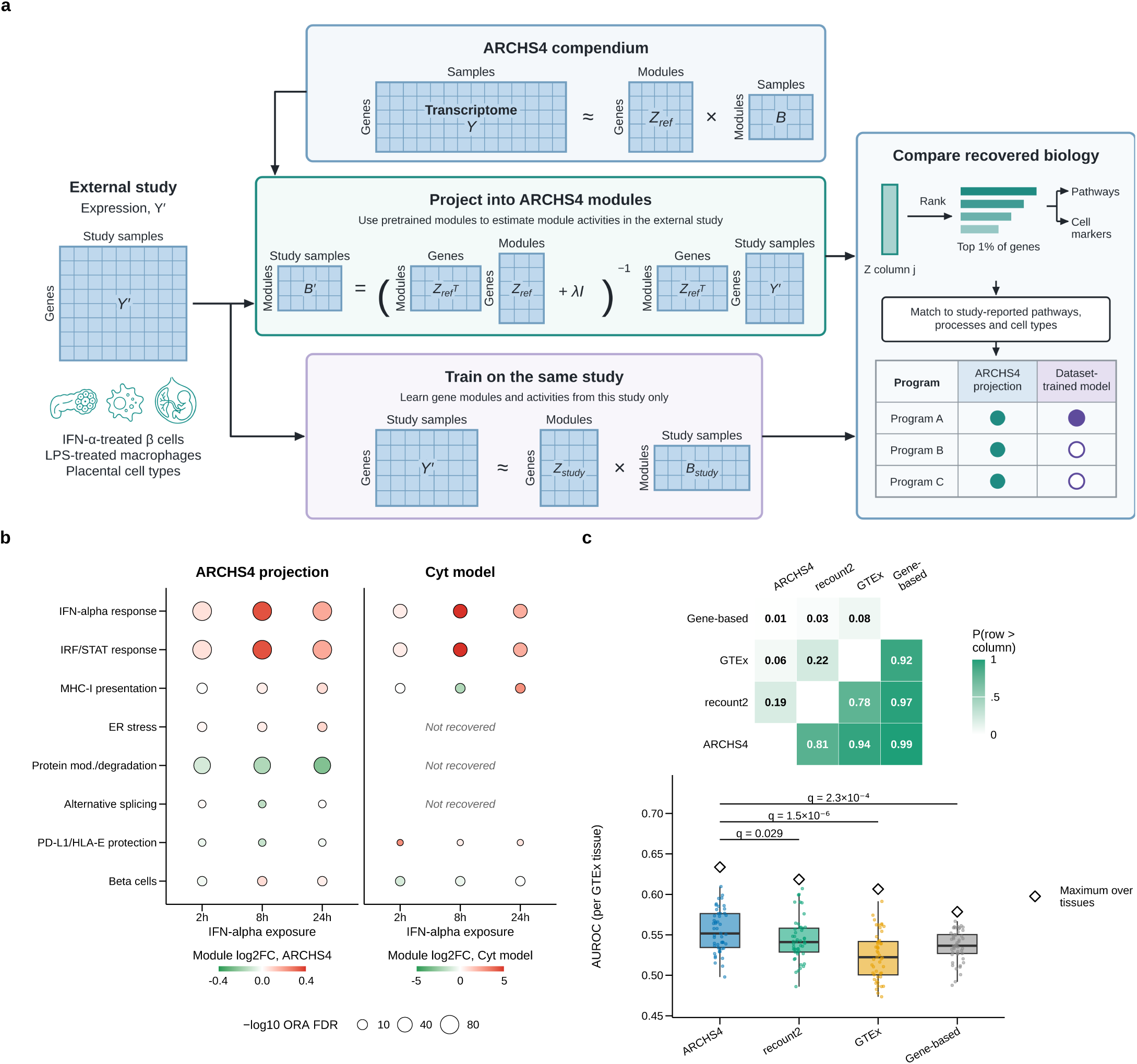
Projection into the ARCHS4 model and drug-disease prediction. a, Projection and comparison design. The ARCHS4 compendium is factorized once into gene loadings *Z_ref_* and module activities *B*. An external study’s expression matrix *Y ′* is either projected into the ARCHS4 modules, estimating activities *B′* with *Zref* fixed by ridge regression with penalty ***λ***, or used to train a CLAMPfull model on the same study (*Zstudy*, *Bstudy*). For each model, the top 1% of genes of each module are tested for enrichment of pathways and cell markers and matched to pathways, cellular processes and cell types reported by the source study. b, IFN-alpha-treated EndoC-betaH1 cells projected into the ARCHS4 model or modelled by a CLAMPfull model trained on the dataset itself (Cyt model). Rows are eight pathways, processes and cell types curated from the source study; fill, module log2 fold change against control at 2, 8 and 24 h (green, decreased; red, increased) on a scale specific to each model; size, -log10(ORA FDR) of the matched gene set; *Not recovered*, no module enriched for a matched gene set at ORA FDR <0.05. c, Drug-disease prediction over 685 labelled drug-disease pairs scored in three CLAMP models and a gene-based approach. Left, fill and value, the fraction of 10,000 paired bootstrap resamples in which the row method reached a higher area under the receiver operating characteristic curve (AUROC) than the column method. Right, AUROC computed separately within each of the 49 GTEx tissues; points, tissues; boxes, medians and interquartile ranges; whiskers, observations within 1.5 interquartile ranges; open diamonds, AUROC of the maximum-over-tissues score; brackets, Benjamini-Hochberg-adjusted paired one-sided Wilcoxon signed-rank tests against ARCHS4 (*q*). IFN, interferon; IRF, interferon regulatory factor; STAT, signal transducer and activator of transcription; ER, endoplasmic reticulum; MHC-I, major histocompatibility complex class I; PD-L1, programmed death-ligand 1; HLA-E, human leukocyte antigen E; ORA, over-representation analysis; FDR, false discovery rate.

We first confirmed that projection preserved known cell-type and tissue signals using the pseudobulk and GTEx benchmarks described above. Because the ARCHS4 representation contains more than an order of magnitude more gene modules than the dataset-trained pseudobulk models (1,728 versus 18-130 per cohort), we measured pseudobulk recovery as the maximum Pearson correlation adjusted by permuting cell-type proportions. In GTEx, we compared the representations at equal module number by clustering samples using 578 of the 1,728 projected module activities, the number of modules in the dataset-trained GTEx models, selected at random in each clustering repetition. In both benchmarks, a null projection was obtained by shu ing entire ARCHS4 gene-loading rows across genes (Methods).

In pseudobulk, projection achieved a mean adjusted maximum Pearson r of 0.552 across 47 dataset-cell-type targets, exceeding the null projection (0.438; q = 1.3 × 10−8; Supplementary Figure 6a). It performed similarly to dataset-trained PLIER (0.552; q = 0.779), with a lower mean than CLAMPbase (0.586; q = 0.088) and CLAMPfull (0.596; q = 0.043). In GTEx, projection recovered subtissue identity with a mean adjusted Rand index (ARI) of 0.642, comparable to dataset-trained CLAMPfull (0.634;q = 0.063) and above PLIER (0.621; q = 4.6 × 10−5), CLAMPbase (0.618; q = 3.9 × 10−4), the expression baseline (0.606; q = 7.7 × 10−11) and the null projection (0.588; q = 2.7 × 10−16; Supplementary Figure 6b). Thus, projection into ARCHS4 retained cell-type and tissue signals comparable to those of models trained on each dataset.

We then asked whether projection recovered biological programs missed by models trained on individual studies, using three published datasets with experimentally characterized responses: EndoC-betaH1 beta cells treated with IFN-alpha for 2, 8 and 24 h^38^, human monocyte-derived macrophages stimulated with LPS^39^ and single-cell RNA-seq of first-trimester placenta, aggregated by donor and cell type^40^. From each publication, we listed the pathways and cellular processes reported by the authors, together with the cell types they described, giving 33 targets: 23 pathways or processes and 10 cell types (Supplementary Table 2). These included IRF/STAT signalling, endoplasmic reticulum (ER) stress and proteasome-mediated protein degradation in beta cells; NAD+ depletion, oxidative stress and a shift towards glycolysis in macrophages; and extravillous trophoblast (EVT) invasion and decidual natural killer (dNK) cell signalling in placenta. Most were also supported in the source studies by measurements other than RNA, such as proteomics, metabolomics, immunohistochemistry and flow cytometry.

For each study, we compared projection into ARCHS4 with a CLAMPfull model trained on the study’s expression data. A target was considered recovered when the top 1% of a module’s gene loadings was enriched for a gene set matched to that target (over-representation analysis (ORA) FDR <0.05; Methods). Projection recovered 20 of 23 pathways or processes and 7 of 10 cell types, compared with 10 and 2 for the dataset-trained models (Figure 4b; Supplementary Figure 6c,d).

Projection also captured more complete programs. Of the nine gene sets recovered by both approaches, ARCHS4 modules included more gene-set members among their top-loading genes for seven (mean 33% versus 22%) (Supplementary Figure 6c-e). The difference was significant for three of them: the interferon-alpha response in beta cells (66% versus 35%; q = 3.8 × 10−5), oxidative phosphorylation in macrophages (55% versus 20%; q = 5.7 × 10−8) and macrophage markers in placenta (15% versus 7%; q = 0.008).

Together, these results show that projection into ARCHS4 recovers pathways, cellular processes and cell types missed by models trained on individual RNA-seq studies, including programs supported by other molecular measurements, and captures more of their genes.

## Larger training compendia improve recovery of known drug-disease relationships

Training on ARCHS4 recovered more pathways, traits and study-reported mechanisms than training on smaller compendia. We next asked whether this advantage extends to drug repurposing approaches that link disease-associated genetic variants to the transcriptional effects of drugs through genetically predicted gene expression^15,41^. These approaches prioritize a drug for a disease when the gene expression changes induced by the drug are opposite to those associated with genetic risk for the disease^41^. Disease-associated expression changes were obtained from S-PrediXcan TWAS results in PhenomeXcan^37,42^, which impute gene expression with prediction models trained in each of 49 GTEx tissues and are based on UK Biobank genome-wide association studies, and drug-induced changes from LINCS L1000 signatures^43^. We compared two implementations of this strategy (Methods): a single-gene approach that scores the opposition between disease and drug profiles across individual genes^41^, and a module-based approach that first projects both profiles into gene modules, as in PhenoPLIER^15^. We applied the module-based approach with the ARCHS4, recount2 and GTEx models and evaluated all four methods on the same manually curated gold standard set of drug-disease medical indications^44^ (Methods).

We first determined how well each method predicted known treatment-disease relationships. We scored drug-disease pairs in each tissue and kept each pair’s best score across the 49 tissues. ARCHS4 achieved the highest AUROC (0.634), followed by recount2 (0.619), GTEx (0.607) and the gene-based approach (0.578; Figure 4c). In 10,000 paired bootstrap resamples, ARCHS4 reached a higher AUROC than the gene-based approach in 99.5% of resamples, than GTEx in 94.1% and than recount2 in 80.7% (Figure 4c). The ordering was unchanged when ranks were averaged over several numbers of top modules or genes (AUROC 0.634 for ARCHS4, 0.613 for recount2, 0.607 for GTEx and 0.581 for the gene-based approach; Supplementary Figure 7a-c).

Because the aggregate score takes a maximum over 49 tissues, its advantage could arise from a few tissues. We therefore compared methods within each tissue. ARCHS4 had the highest median tissue AUROC (0.552) and was significantly better than recount2 (0.541; q = 0.029), the gene-based approach (0.537; q = 2.3 × 10−4) and GTEx (0.522; q = 1.5 × 10−6; Figure 4c; Supplementary Figure 7d).

Together, these results show that gene modules learned from larger compendia improve drug-disease prediction over both single genes and modules learned from smaller compendia, across and within tissues.

## Consensus gene modules connect CRISPR-validated lipid genes missed by single-gene TWAS to disease programs

Finally, we asked whether ARCHS4 modules, integrated with TWAS results and drug-perturbation signatures, recover disease-relevant programs that include experimentally validated genes. We reanalysed the two high-confidence gene sets, lipid-increasing and lipid-decreasing, of a fluorescence-based CRISPR-Cas9 screen in HepG2 cells^15^ (Methods). Because matrix factorization is stochastic, independent runs can converge to different solutions, so a module learned in a single run may be an artefact of that run rather than a robust biological signal. Modules can also capture technical factors, such as batch effects, rather than biology. We therefore trained, in addition to the ARCHS4 model used in the previous analyses, nine CLAMPfull models on ARCHS4 in exactly the same way but with different random initializations, and removed modules associated with gene-or sample-level technical covariates from all ten models (Methods; Supplementary Table 3). To retain only reproducible programs, we clustered the remaining modules by the cosine similarity of their gene loadings (complete linkage, similarity ≥0.5); 618 of the 8,025 clusters contained modules from at least six of the ten models, and we combined the modules of each into a consensus meta-gene module (MGM) by taking, for every gene, the median loading across them.

We then tested the MGMs for enrichment of the lipid-increasing and lipid-decreasing CRISPR gene sets. Twenty-seven of the 618 MGMs were enriched for at least one set (adjusted P <0.05; Supplementary Figure 8; Supplementary Table 4). For each enriched MGM, we used its top 1% of genes by loading to compute genetic trait associations with PhenoPLIER (Supplementary Table 5), canonical pathway and cell-type marker enrichment (Supplementary Table 6) and the response of these genes to drug perturbations in LINCS L1000 expression profiles (Supplementary Table 7). We also characterized the tissue and cell-type context of each MGM by annotating the 1% of ARCHS4 samples with the highest MGM activity (Methods). Four MGMs were associated with a trait (FDR <0.05) and enriched for a lipid-related canonical pathway (FDR <0.05) (Figure 5). MGM91 was reproduced in all ten models, MGM175 in nine, MGM335 in seven and MGM588 in six, with a mean pairwise cosine similarity between member modules of 0.81, 0.75, 0.75 and 0.85, respectively.

**Figure 5:**
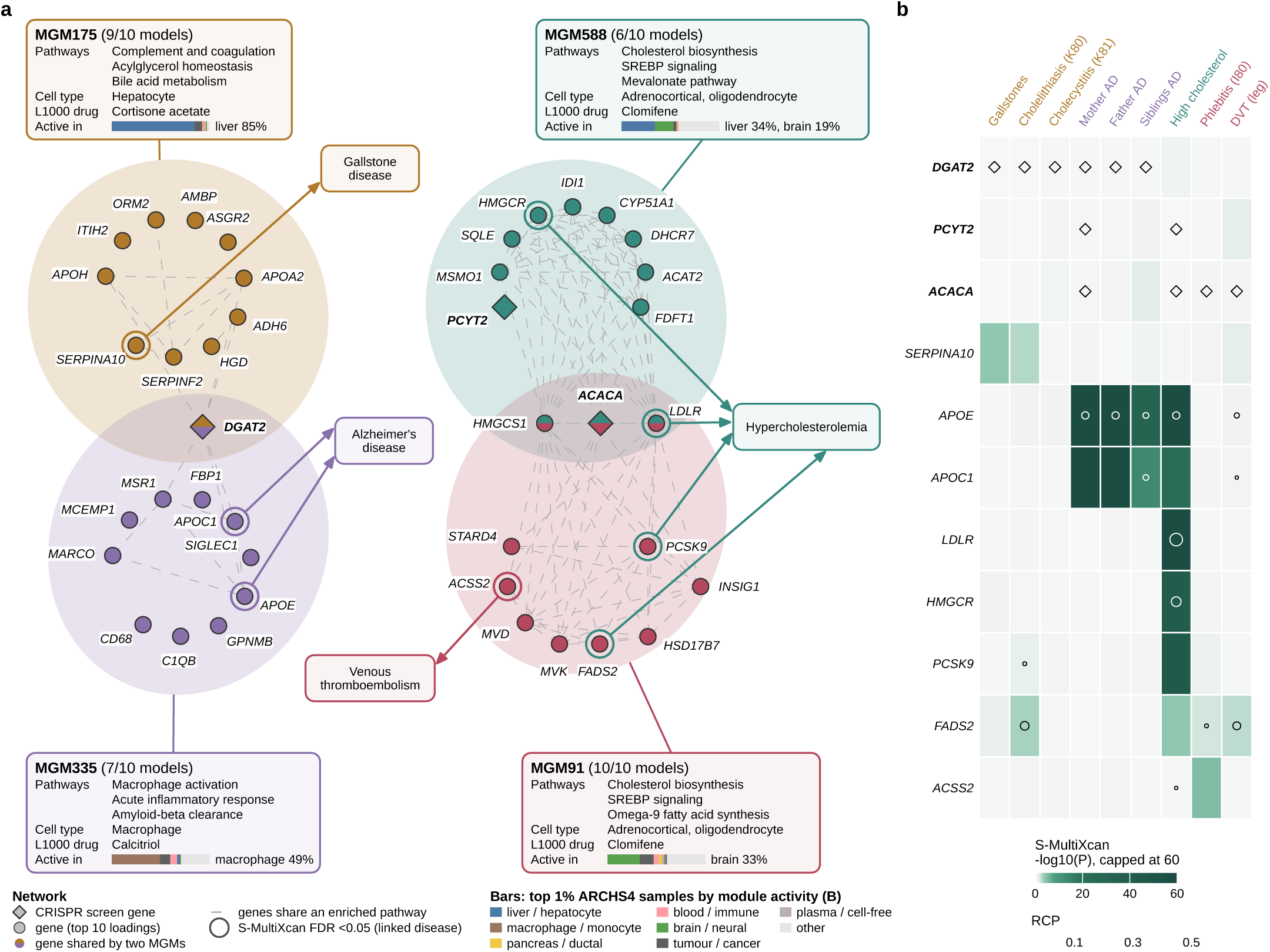
Lipid-screen MGMs connect experimentally validated genes to disease. a, The four MGMs, each reproduced in at least six of ten CLAMPfull models trained on ARCHS4, enriched for a high-confidence HepG2 lipid-screen gene set (fgsea adjusted *P* <0.05), associated with a trait at false discovery rate (FDR) <0.05 and enriched for a lipid-related canonical pathway (Supplementary Tables 4, 5 and 6). Each shaded circle is one MGM, showing its CRISPR-Cas9 screen genes (diamonds) and its top 10 genes by loading (dots); genes shared by two MGMs lie in the overlap of their circles and are coloured by both. Grey dashed edges, gene pairs within an MGM that share a pathway enriched in that MGM (ORA FDR <0.05); ringed genes, S-MultiXcan FDR <0.05 for the linked disease; arrows, MGM-disease links, merging traits at GLS FDR <0.05 with at least 1,000 cases into four diseases. Boxes give the number of models reproducing the MGM, its top three canonical and GO Biological Process pathways, its cell type from CellMarker and Azimuth marker enrichment, the LINCS L1000 drug ranked highest among those acting on the program’s biology (Supplementary Table 7) and the sample types of the 1% of ARCHS4 samples with the highest MGM activity (labelled when ≥15%). b, S-MultiXcan transcriptome-wide associations in PhenomeXcan for the three screen genes (bold) and reference genes across the traits associated with the MGMs in a (trait labels coloured by disease). Fill, -log10(*P*) capped at 60; circles, fastENLOC regional colocalization probability (RCP) ≥0.01, sized by RCP; diamonds, each screen gene with the traits associated with the MGMs that carry it. AD, Alzheimer’s disease; DVT, deep venous thrombosis; MGM, meta-gene module; ORA, over-representation analysis; TWAS, transcriptome-wide association study.

For each of the four MGMs, these independent data sources, pathways, drug responses and genetic trait associations together with cell-type markers and the most active samples, converged on the same biological process (Figure 5a). For example, MGM175, a hepatocyte program led by *APOH*, *ITIH2*, *ORM2* and *AMBP* and most active in liver samples, was enriched for complement and coagulation, acylglycerol homeostasis and bile acid metabolism, was induced by the glucocorticoid cortisone acetate and was associated with gallstone disease (Supplementary Table 5). The other three MGMs showed the same pattern (Figure 5a). MGM335, a macrophage program (*MARCO*, *MSR1* and *APOE*) involved in amyloid-beta clearance, was induced by calcitriol, the active form of vitamin D, and was linked to family history of Alzheimer’s disease. MGM91 combined cholesterol and fatty-acid synthesis, whereas MGM588 centred on the mevalonate pathway; both were induced by clomifene, an inhibitor of sterol synthesis, and were linked to self-reported high cholesterol, but MGM91 was also linked to venous thromboembolism (Supplementary Table 5).

None of the screen genes that drove the enrichment of these MGMs, *ACACA*, *DGAT2* and *PCYT2*, was detected by single-gene TWAS for the traits associated with their programs (Figure 5b; Supplementary Table 8): the smallest *P* was 0.21, for *ACACA* and phlebitis, and none showed evidence of colocalization between GWAS and expression quantitative trait locus signals (regional colocalization probability (RCP) ≤0.004). The function of each screen gene matched its program: *DGAT2*, which catalyses the final step of triglyceride synthesis, lay in the hepatocyte program enriched for acylglycerol homeostasis (MGM175) and in the macrophage program enriched for lipid signalling (MGM335); *ACACA*, the first enzyme of fatty-acid synthesis, lay in MGM91, which combined cholesterol and fatty-acid synthesis, and in MGM588; and *PCYT2*, the rate-limiting enzyme of phosphatidylethanolamine synthesis, lay in the sterol-synthesis program MGM588. By contrast, other genes of the same programs were strongly associated with the linked diseases, including *SERPINA10* with gallstones (P = 7.8 × 10−5), *LDLR* (P = 9.9 × 10−136) and *PCSK9* (P = 1.3 × 10−45) with high cholesterol, and *ACSS2* with venous thromboembolism (P = 3.3 × 10−7).

Together, MGMs embedded experimentally validated lipid-regulating genes in mechanistically coherent programs whose genes, cellular contexts and drug responses converged on disease associations, linking these genes to disease through the programs that contain them even when single-gene TWAS did not detect them.

## Discussion

Interpreting genetic risk through gene programs requires methods that accurately resolve cell- and tissue-specific programs and that can be trained on expression data diverse enough to contain them. Here we present CLAMP, a scalable prior-informed matrix factorization framework that learns interpretable gene modules from large expression compendia. CLAMP recovered cell-type composition and tissue organization more accurately than state-of-the-art decomposition methods, and retained cell-type signals in bulk expression data, where cellular composition is mixed. Its scalability enabled training on virtually all public human bulk RNA-seq in ARCHS4. Modules learned from this larger compendium recovered more pathways and genetic traits than those learned from smaller compendia and, through projection, recovered programs missed by models trained on individual studies. When used to connect genetic associations with drug-induced expression changes, these modules ranked known drug-disease pairs better than modules learned from smaller compendia or individual genes. Integrated with TWAS and drug-perturbation data, they linked CRISPR-validated lipid regulators missed by single-gene TWAS to disease.

CLAMP differs from previous prior-informed methods in how it uses curated gene sets. PLIER encourages gene loadings to resemble weighted combinations of pathways, and MOFA-FLEX links each factor to a predefined gene set^11,12^. CLAMP instead identifies the gene sets associated with each module learned from the expression data and uses them to adjust how strongly each gene’s loading is shrunk: genes supported by these gene sets are shrunk less toward zero, and all other genes more strongly. Modules can therefore align with known pathways while still capturing co-expressed genes not yet annotated to them. When trained with the same gene sets, CLAMP recovered cell-type and tissue signals more accurately than PLIER and MOFA-FLEX. In addition, algorithmic optimizations and file-backed computation reduced runtime and memory requirements relative to other prior-informed methods (Supplementary Figure 4), which made it possible to train on more than 600,000 samples and to build the ARCHS4 reference model.

Beyond benchmark accuracy, CLAMP modules allowed independent lines of evidence to be evaluated on the same biological processes. Interpreting individual latent factors is risky because matrix factorization depends on initialization and can capture technical variation^11,45^. We therefore restricted interpretation to consensus modules (MGMs) reproduced across ten independently trained models after removing modules associated with technical covariates. Within these MGMs, cell-type identity, pathway activity, drug responses and genetic associations, each derived from independent data, converged on the same process. This convergence provides internal support for module interpretation in the absence of ground truth, and it was reflected at the gene level: the function of each screen gene matched its program, as for *DGAT2*, which catalyses the final step of triglyceride synthesis, in a hepatocyte program enriched for acylglycerol homeostasis, *ACACA*, the first enzyme of fatty-acid synthesis, in a program combining cholesterol and fatty-acid synthesis, and *PCYT2*, the phosphatidylethanolamine synthesis enzyme, in a sterol-synthesis program. None of the screen genes had a detectable TWAS association or colocalization signal of its own, yet each was linked to disease through other gene members of its programs with strong associations, such as *LDLR* and *PCSK9* for high cholesterol. Extending earlier module-based analyses of this screen^15^, this pattern is consistent with omnigenic and stratagenic models, in which genes with direct functional effects may lack detectable individual associations while their programs carry the genetic signal^1,3^. The same lipid processes also connected metabolic, hepatic, vascular and neurodegenerative traits, including high cholesterol, gallstone disease, venous thromboembolism and Alzheimer’s disease, consistent with network pleiotropy, in which shared programs mediate molecularly related traits. By connecting genetic, transcriptional and perturbational evidence through shared programs, CLAMP thus provides a way to operationalize omnigenic and stratagenic models of complex disease, prioritize candidate genes that single-gene approaches miss and generate testable hypotheses about the cell types and processes through which genetic risk acts.

This study has limitations. First, CLAMP was trained on bulk expression, which averages signals across cells and can therefore mask programs of rare cell types or transient cell states. CLAMP partly overcame this limitation, as modules trained on bulk-like pseudobulk recovered cell-type proportions in held-out donors and identified the corresponding cells when single cells were projected onto them. Because its optimized, file-backed training scales to hundreds of thousands of samples, CLAMP could be applied to large single-cell compendia to resolve cell types and cell states at higher resolution. Second, the stratagenic model proposes that epigenetic and environmental factors converge on the same programs as genetic risk^3^, whereas CLAMP currently connects programs only to genetic associations and drug perturbations; projecting chromatin accessibility, DNA methylation and environmental exposures into the module space could reveal how these factors jointly shape disease-relevant programs. Third, our genetic analyses relied on cis-based TWAS, which does not capture trans-regulatory effects, and integrating trans-eQTLs could identify trans regulators of disease-associated programs. Finally, module-level analysis prioritizes candidate genes but does not establish causality: the genes highlighted here had experimental support from a single CRISPR screen of lipid accumulation in HepG2 cells, and determining whether candidate genes causally affect disease will require targeted perturbations in the cell types indicated by each program.

Overall, CLAMP provides a framework for interpreting genetic associations through cellular programs learned from public RNA-seq data, supporting the functional characterization of complex disease.

CLAMP is available as an open-source R package through Bioconductor, together with what is, to our knowledge, the largest interpretable gene-module model of human bulk RNA-seq to date, trained on 605,614 ARCHS4 samples, and the GTEx and recount2 models, released as a browsable atlas annotated with pathways, cell types, drug perturbations and genetic traits. New expression datasets can be projected into this reference without retraining, and new TWAS results and drug signatures can be analysed against its modules with the PhenoPLIER framework, so that programs learned from public RNA-seq data can be reused across studies and extended as these data continue to grow.

## Methods

### CLAMP algorithm

CLAMP approximates a standardized gene-by-sample expression matrix Y as Y ≈ ZB, where Z (genes × modules) contains non-negative gene loadings and B (modules × samples) contains module activities. The rank k was estimated from the singular-value spectrum of Y , computed by randomized singular value decomposition. B was initialized from the leading right singular vectors, and the penalties were set to λ1 = dk/3 and λ2 = dk, where d_k_ is the *k*th singular value.

*CLAMPbase* fits the model without prior information:

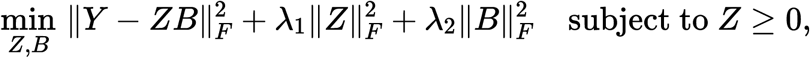

by alternating ridge updates of *Z* and *B*. After each *Z* update, negative loadings and small positive loadings (below the magnitude of the 5% quantile of the module’s negative loadings) were set to zero. Fitting stopped when the relative change in *B* fell below 5 × 10−4.

*CLAMPfull* is initialized from CLAMPbase and adds a binary gene-by-gene-set prior matrix *C*. For each module, the loadings were regressed on the 10 best-discriminating gene sets (ranked by AUROC) by non-negative elastic-net regression (glmnet^46^, α = 0.9), with the penalty selected by five-fold cross-validation. The selected sets were then refitted without penalty, giving the coefficient matrix *U*. The predicted loadings *CU* define a multiplier *mgj* ≥ 0, which is the excess of the predicted loading variance of gene *g* over the background loading variance of module *j. Z* and *B* are then estimated as

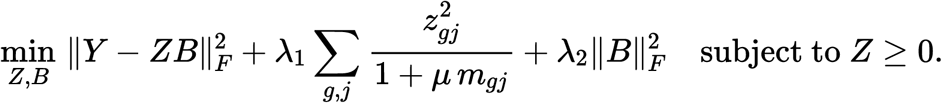

Genes supported by the selected gene sets are therefore shrunk less towards zero, while genes without prior support keep the base penalty and can still carry large loadings. The constant *μ* sets the strength of the prior (100 for the reference models). *U* was updated periodically during training. To validate module-gene-set associations, 20% of the genes in each gene set were withheld from *C* during training, and their loadings were scored by AUROC with a rank-sum test and BH correction.

*Projection.* New data *Y ′* were projected without refitting the loadings as *B′ = (Z ⊤Z + λ2I)−1Z ⊤Y ′* . Large compendia were stored as file-backed matrices (bigstatsr^47^) and processed blockwise.

### Pseudobulk generation

Two types of donor-level profiles were generated from six single-cell cohorts. For pseudobulk profiles, counts were normalized to counts per million (CPM) within each cell and averaged across all annotated cells from each donor, so that every cell contributed equally regardless of sequencing depth. Duplicate gene symbols were averaged, and donors with fewer than 400 cells were excluded. For overall pseudobulk profiles, raw counts were summed across all annotated cells from each donor, regardless of cell type and without per-cell normalization, and then converted to CPM.

### Datasets and preprocessing

The single-cell cohorts were Mathys et al.^19^, Xiong et al.^20^, HeartMap^22^, the Human Lung Cell Atlas^21^, OneK1K^17^ and Perez et al.^18^. The bulk compendia were GTEx v8^4^, ARCHS4 v2.5^16^ and recount2^48^ as processed for MultiPLIER^13^. Bulk TPM (GTEx, ARCHS4) and RPKM (recount2) were log2(x + 1)-transformed, whereas pseudobulk CPM were used on the linear scale. ARCHS4 libraries with a single-cell probability ≥0.5 were removed. In all datasets, genes with mean >0.5 and variance >0.1 were retained and standardized to zero mean and unit variance. After preprocessing, GTEx comprised 21,613 genes × 17,382 samples, ARCHS4 18,423 × 605,614 and recount2 5,993 × 37,032.

### Pseudobulk benchmarking

CLAMPbase and CLAMPfull were compared with PCA^25^, ICA^27^, NMF^26^, PLIER^11^, MOFA-FLEX^12^, flashier^23^, GenomicSuperSignature^14^ and CoGAPS^24^ on 47 dataset-cell-type targets. Prior-informed methods received the same 5,343 Gene Ontology Biological Process terms^49^, obtained from Enrichr (2025)^50^. No cell-type markers were used for training. The rank was twice the elbow of the singular-value spectrum. Cell types were matched one-to-one to modules by greedy maximum Pearson correlation between proportion and activity, and recovery was the correlation of the matched module. Generalization was assessed by five-fold donor-grouped cross-validation, with all preprocessing, rank selection, fitting and assignment estimated on training donors only; a univariate least-squares regression fitted on training donors converted the assigned module’s activity into a predicted proportion. Module specificity was summarized by Tau^51^ and donor-level AUROC. Module genes (the top 1% of loadings) were annotated by over-representation analysis (ORA; one-sided hypergeometric test; clusterProfiler^52^) against CellMarker 2.0^29^, Azimuth^28^ and Allen Brain Atlas^30^ markers. Single cells were projected into the fitted models without refitting. Overall pseudobulk profiles were analysed in the same way as the pseudobulk profiles.

### GTEx analyses

The GTEx rank was twice the Gavish-Donoho estimate^35^ (578 modules). CoGAPS did not converge within seven days and was excluded. Sample representations were clustered by *k*-means (*k* = 54 subtissues; 50 seeded repetitions), and agreement with subtissue labels was measured by adjusted Rand index (ARI). Methods were compared by paired one-sided Wilcoxon signed-rank tests. The expression baseline used the 25% most variable genes. For subtissues with ≥50 samples, one-versus-rest random forests with TreeSHAP^53^ were used to rank module importance, and the top modules were annotated by ORA against 511 GTEx tissue signatures. Subtissue prediction in held-out donors used multinomial logistic regression with donor-grouped cross-validation, permutation tests and donor bootstraps. Cell-type recovery was compared with xCell scores^33^ from Kim-Hellmuth et al.^34^.

### Reference models

The ARCHS4 (1,728 modules), GTEx (578) and recount2 (544) reference models were CLAMPfull fits with a combined prior of MSigDB C2-all^36^, Reactome^54^ and CellMarker (release 2026.1.Hs). They were used for projection, drug-disease prediction and the CRISPR analyses.

### Training-size and rank analyses

ARCHS4 was subsampled by whole studies to 1, 5, 10, 25, 50 and 75% of samples (three seeds each), and three independently seeded fits were used at 100%. Each subset used the same genes and preprocessing statistics as the full compendium, so models differed only in their training samples, and the number of modules was re-estimated for each subset. Additional models were fitted at fixed ranks from 86 to 1,728; one 1% subsample yielded fewer singular components (1,527) than 1,728 modules and was omitted at that rank. Pathway coverage was the fraction of MSigDB canonical pathways (excluding Reactome)^36^ and CellMarker cell-type gene sets^29^ (10-500 genes) recovered by ORA (BH-adjusted P <0.05) in at least one CLAMPbase module. Genetic-trait recovery used PhenoPLIER generalized least squares (GLS)^15^ on PhenomeXcan associations^37^ for 2,366 traits. A trait was considered recovered if at least one module had FDR <0.05.

### Projection of external datasets

External datasets were projected into the ARCHS4 reference on shared genes. In the pseudobulk transfer comparison, each correlation was reduced by the median of a null distribution from 10,000 proportion permutations, to account for the larger number of modules searched. Null projections permuted gene identities in the loading matrix. Reference projection was compared with locally fitted models in IFN-α-treated EndoC-βH1 cells (GSE133218)^38^, LPS-treated macrophages (GSE193336) and placental cells (E-MTAB-6701)^40^. Differential module activity was tested with limma^55^, with donor included as a covariate for datasets with several samples per donor (BH-adjusted P <0.05).

### Drug-disease prediction

As in PhenoPLIER^15^, LINCS L1000 drug signatures^43^ and S-PrediXcan z-scores for 49 tissues were projected into each reference model. For each trait, the top 5, 10, 25 or 50 modules by absolute association were retained, and the drug-trait score was the negative dot product of the two vectors. The single-gene baseline applied the same rule to the top 50-500 genes. Traits were mapped through the Experimental Factor Ontology^56^ to Disease Ontology^57^ terms, and scores were averaged across thresholds and maximized across tissues. Predictions were evaluated against curated indications^44^ on 685 pairs (531 positive and 154 negative) by AUROC and average precision. Differences were tested with 10,000 paired bootstrap resamples.

### Reanalysis of the CRISPR lipid screen

The two high-confidence gene sets from a published HepG2 CRISPR-Cas9 lipid screen^15^ were reanalysed: six lipid-increasing and eight lipid-decreasing genes. Meta-gene modules (MGMs) were built from ten independently seeded ARCHS4 reference models. Modules were first flagged as technical and removed if their loadings correlated with transcript length or GC content, or if technical covariates explained ≥25% of their activity variance (cross-validated R²). The remaining loadings were clustered by complete linkage on cosine distance (cut at similarity 0.5). Clusters containing modules from at least six models were retained (618 MGMs), and each was summarized by the gene-wise median of its members’ loadings. MGMs enriched for either screen set were identified with fgsea^58^ (adjusted P <0.05), then tested for trait associations with PhenoPLIER GLS, annotated by ORA and scored against 1,170 LINCS drug signatures. For sample context, MGM loadings were projected onto all ARCHS4 samples, and the 1% of samples with the highest activity were annotated from their GEO source, title and characteristics by keyword rules.

### Computational performance

Runtime and memory were measured on an AMD Ryzen Threadripper PRO 5955WX workstation (16 cores, 128 GB RAM) with four CPU threads, GPUs disabled and seeds 123, 456 and 789. Wall-clock times included input loading, model fitting and serialization, and peak GTEx memory was the maximum resident set size.

## Data availability

All data analysed are publicly available from the sources cited above. The trained CLAMP models for ARCHS4, GTEx and recount2, including gene loadings, sample activities and module annotations, are released through a browsable web application.

## Code availability

The CLAMP R package is available from Bioconductor (https://bioconductor.org/packages/CLAMP) and GitHub (https://github.com/chikinalab/CLAMP). Analysis code is available at https://github.com/pivlab/clamp-analyses. Analyses used CLAMP 0.99.0, R 4.4.3 and Python 3.11.

## Acknowledgements

This work was supported by the National Human Genome Research Institute (R00 HG011898 to M.P. and R01 HG009299-6A1 to M.C.), the Eunice Kennedy Shriver National Institute of Child Health and Human Development (R01 HD109765 to M.P.), the National Science Foundation (2238125 to M.C.), the National Eye Institute (R01 EY030546-01A1 to M.C.) and the National Library of Medicine (T15LM009451 to J.H.).

