## Supplementary Information, Notes, Figures and Tables for "A genetically informed atlas of transcriptomic programs from massive human RNA-seq compendia"

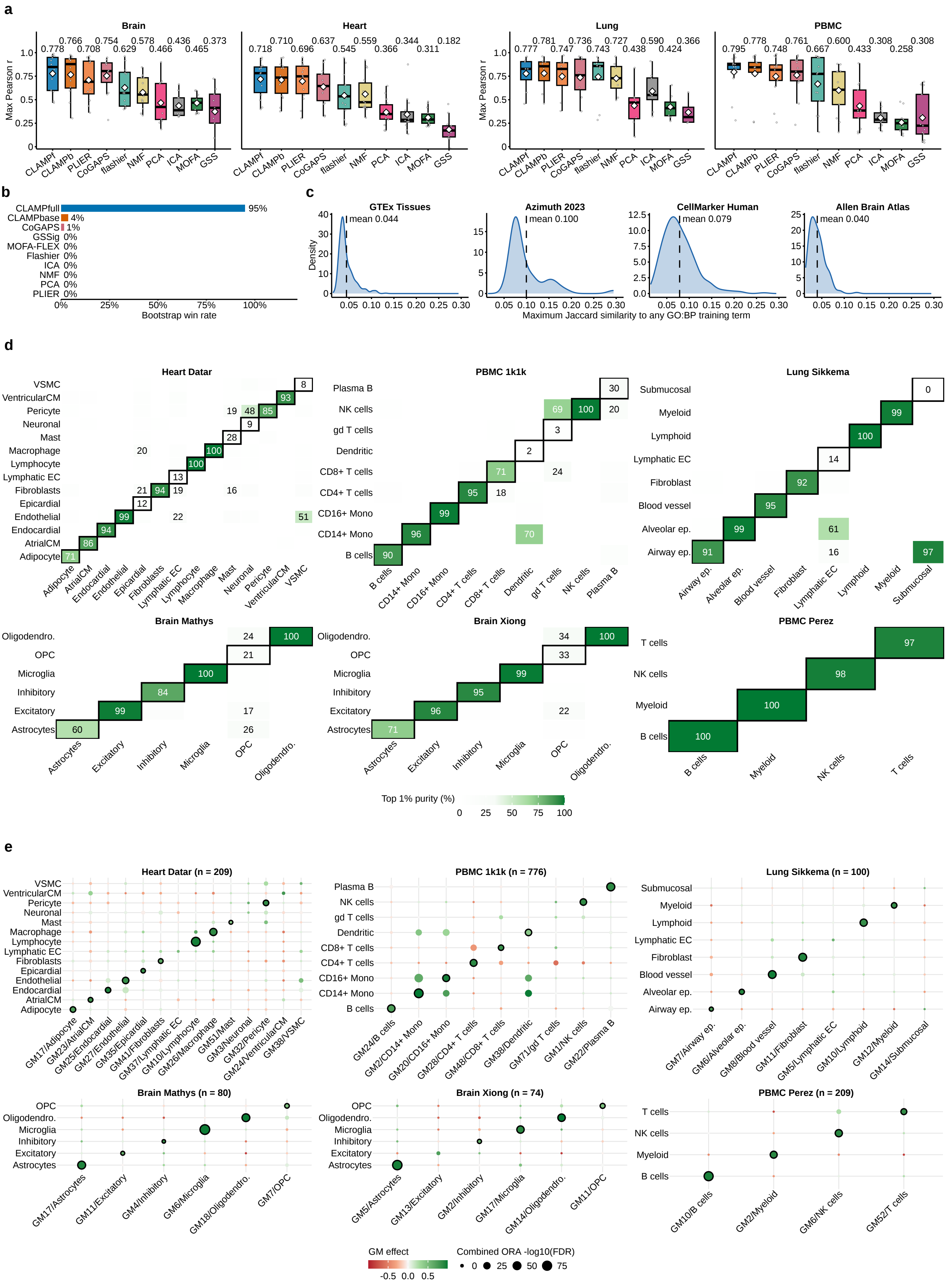

**Supplementary Figure 1: Robustness of pseudobulk benchmarking and cell-type recovery. a**, Benchmark of cell-type proportion recovery across methods, separated by tissue: brain ($n = 12$ targets), heart ($n = 14$), lung ($n = 8$) and PBMC ($n = 13$), quantified by the maximum Pearson correlation as in Figure 2a. Points, targets; boxes, median and interquartile range; whiskers, values within 1.5× the interquartile range; diamonds and labels, means. **b**, Fraction of 2,000 dataset-level bootstrap resamples in which each method achieved the highest mean correlation across targets. **c**, Maximum Jaccard similarity between each evaluation marker set and any of the 5,343 GO:BP terms used as training priors. Distributions are shown for GTEx tissue, Azimuth, CellMarker 2.0 and Allen Brain Atlas gene sets; dashed lines, collection means. **d**, Recovery of cell identities after projection of the original single cells into modules learned from donor-level pseudobulk profiles. Values indicate the percentage of each source cell annotation among the top 1% highest-activity cells for each assigned module. Rows, cell annotations; columns, assigned modules; outlines, matched annotation-module pairs; off-diagonal values are shown when ≥15%. **e**, Cell-type marker enrichment among the top 1% of genes ranked by signed module loading for all 47 assigned modules. Columns, modules labelled as GM/assigned cell type and ordered to match the rows; rows, marker sets. Point size indicates combined ORA $-\log_{10}(\mathrm{FDR})$; colour indicates the correlation between module activity and cell-type proportion; outlines, matched module-cell-type pairs. Matched marker enrichment was recovered for 39 of 47 modules. Dataset titles indicate donor numbers. GO:BP, Gene Ontology Biological Process; ORA, over-representation analysis; FDR, false discovery rate; GM, gene module; OPC, oligodendrocyte precursor cell; EC, endothelial cell; Mono, monocyte; CM, cardiomyocyte; ep., epithelium; VSMC, vascular smooth muscle cell.

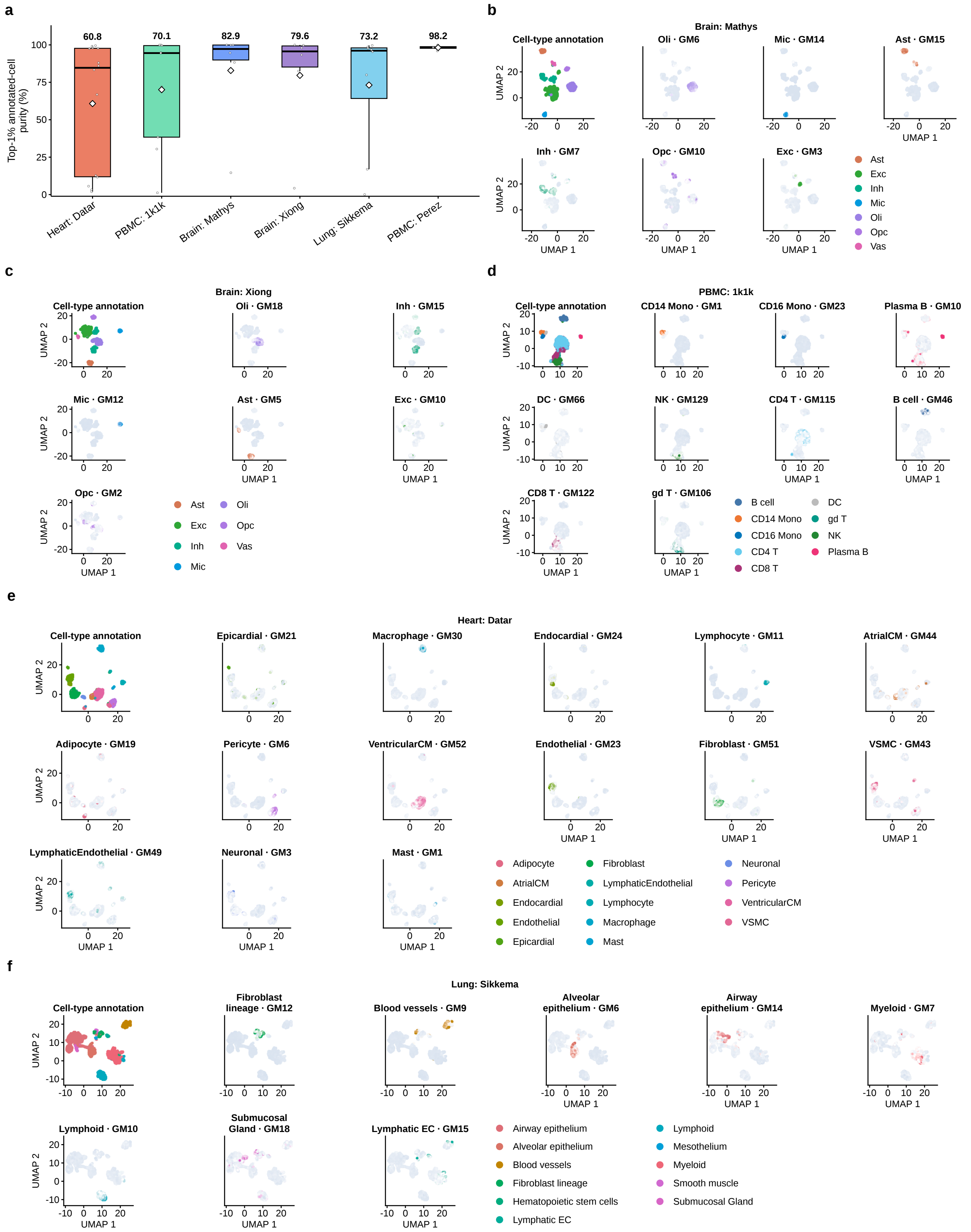


**Supplementary Figure 2: Single-cell recovery from donor-level bulk-like expression profiles. a**, Percentage of cells with the matched annotation among the top 1% highest-activity cells for each assigned module. Results are shown for Heart: Datar ($n = 14$ modules), PBMC: 1k1k ($n = 9$), Brain: Mathys ($n = 6$), Brain: Xiong ($n = 6$), Lung: Sikkema ($n = 8$) and PBMC: Perez ($n = 4$). Points, modules; boxes, median and interquartile range; whiskers, values within 1.5× the interquartile range; diamonds and labels, means. **b–f**, Single-cell projections for Brain: Mathys, Brain: Xiong, PBMC: 1k1k, Heart: Datar and Lung: Sikkema, respectively. For each dataset, the first map shows source cell-type annotations on an expression-derived UMAP computed independently of CLAMP, and the remaining maps show activity of the module assigned to each indicated cell type on the same embedding. Module activities range from pale gray-blue for low activity to the corresponding cell-type colour for high activity. UMAPs show up to 60,000 cells per dataset, subsampled by annotated cell type. GM, gene module; UMAP, uniform manifold approximation and projection; PBMC, peripheral blood mononuclear cell; CM, cardiomyocyte; EC, endothelial cell; VSMC, vascular smooth muscle cell; Mono, monocyte; DC, dendritic cell; NK, natural killer cell; gd T, gamma-delta T cell; Exc, excitatory neuron; Inh, inhibitory neuron; Ast, astrocyte; Oli, oligodendrocyte; OPC, oligodendrocyte precursor cell; Mic, microglia; Vas, vascular cell.

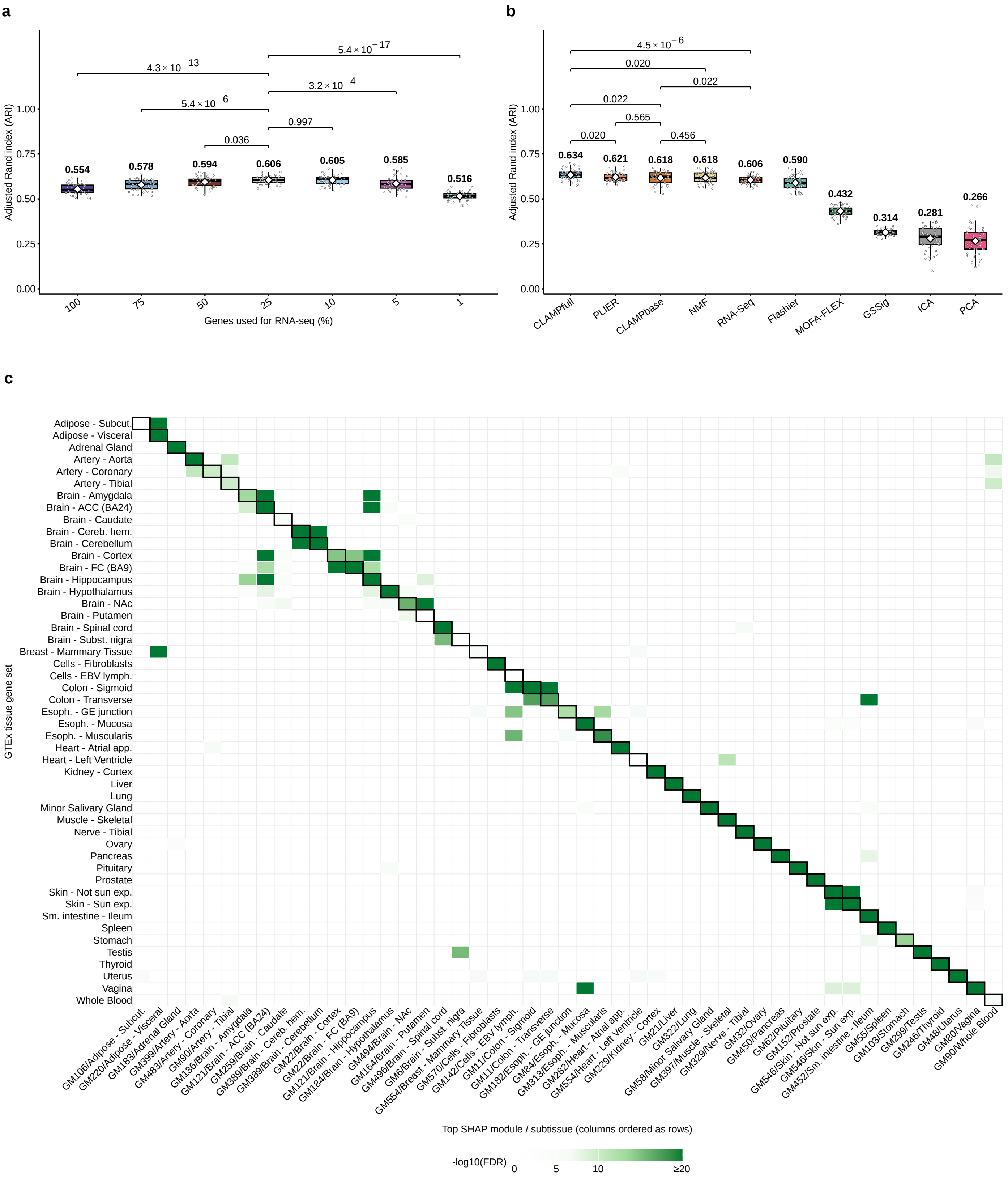


**Supplementary Figure 3: GTEx tissue organization and recovery of tissue-associated gene programs. a**, Sensitivity of GTEx subtissue recovery to the fraction of highly variable genes used for the RNA-seq expression baseline, quantified by adjusted Rand index (ARI) between $k$-means clusters ($k = 54$) and subtissue labels ($n = 50$ repetitions per gene fraction). Points, repetitions; boxes, median and interquartile range; whiskers, values within 1.5× the interquartile range; diamonds and labels, means; brackets, Benjamini-Hochberg-adjusted two-sided Mann-Whitney tests for the indicated comparisons. **b**, Benchmark of subtissue recovery across methods using ARI, with expression restricted to the 25% most variable genes as the RNA-seq baseline ($n = 50$ matched repetitions). Points, boxes, whiskers, diamonds and labels are as in **a**; brackets, Benjamini-Hochberg-adjusted paired one-sided Wilcoxon tests for the indicated comparisons. **c**, Enrichment of GTEx tissue gene sets among the top 1% of gene loadings from the highest SHAP-ranked module for each of 49 subtissues. Rows indicate GTEx tissue gene sets; columns indicate the top-ranked module for each subtissue and are ordered to match the rows. A module ranked first for multiple subtissues therefore appears in multiple columns. Colour indicates $-\log_{10}(\mathrm{FDR})$, capped at 20, for FDR < 0.0001; white, FDR ≥ 0.0001; outlines, expected tissue-module pairs. Each tile reports the minimum FDR across the sex- and age-stratified GTEx signatures for that tissue, with Benjamini-Hochberg correction performed within each module. ARI, adjusted Rand index; SHAP, Shapley additive explanations; FDR, false discovery rate; TPM, transcripts per million; GM, gene module; ACC, anterior cingulate cortex; FC, frontal cortex; NAc, nucleus accumbens; Cereb. hem., cerebellar hemisphere; Subst. nigra, substantia nigra; EBV lymph., EBV-transformed lymphocytes; Esoph., esophagus; GE junction, gastroesophageal junction; Atrial app., atrial appendage; Sm. intestine, small intestine; Subcut., subcutaneous; Sun exp., sun-exposed.

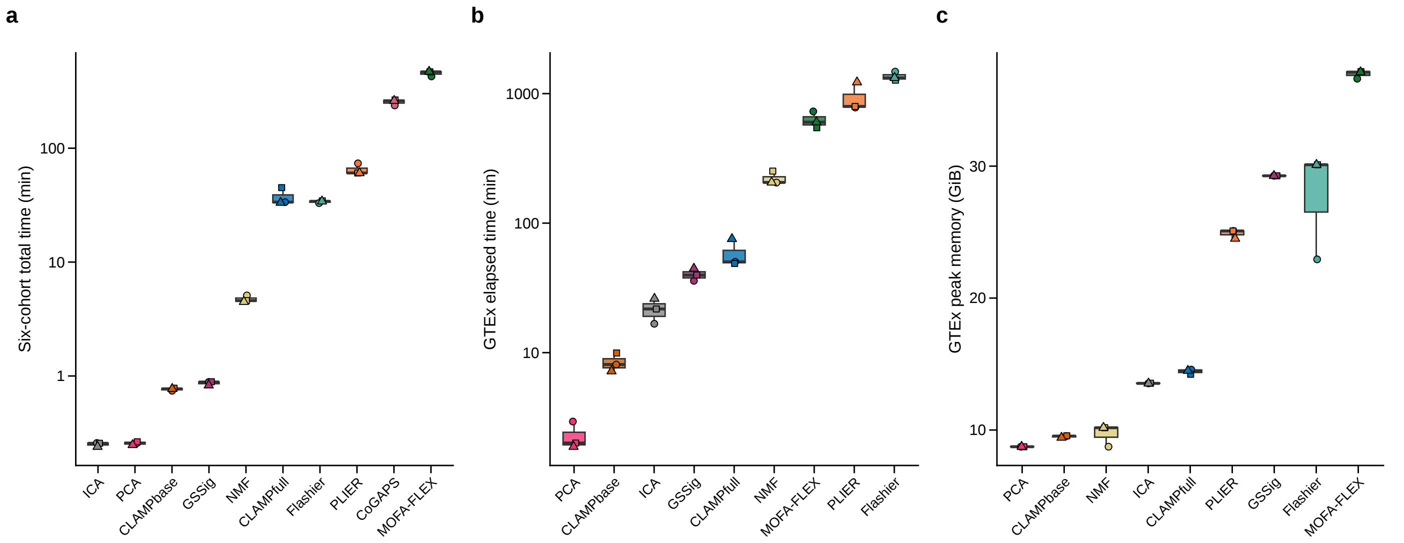


**Supplementary Figure 4: Computational performance across pseudobulk and GTEx analyses. a**, Total wall-clock time required to fit all six pseudobulk datasets, summed within each of three seeds per method. **b**, Wall-clock time required to fit GTEx for three seeds per method. **c**, Peak memory during GTEx model fitting. Points indicate individual seeds (123, 456 and 789), distinguished by shape; boxes, median and interquartile range; whiskers, values within 1.5× the interquartile range. Time axes in **a** and **b** are logarithmic. Timings include input loading, model fitting and serialization, with GTEx input-cache warming performed before measurement. CoGAPS is omitted from **b** and **c** because the GTEx run exceeded seven days without convergence. Memory is reported in binary gigabytes (GiB; $2^{30}$ bytes). Additional measurement details are provided in Methods.

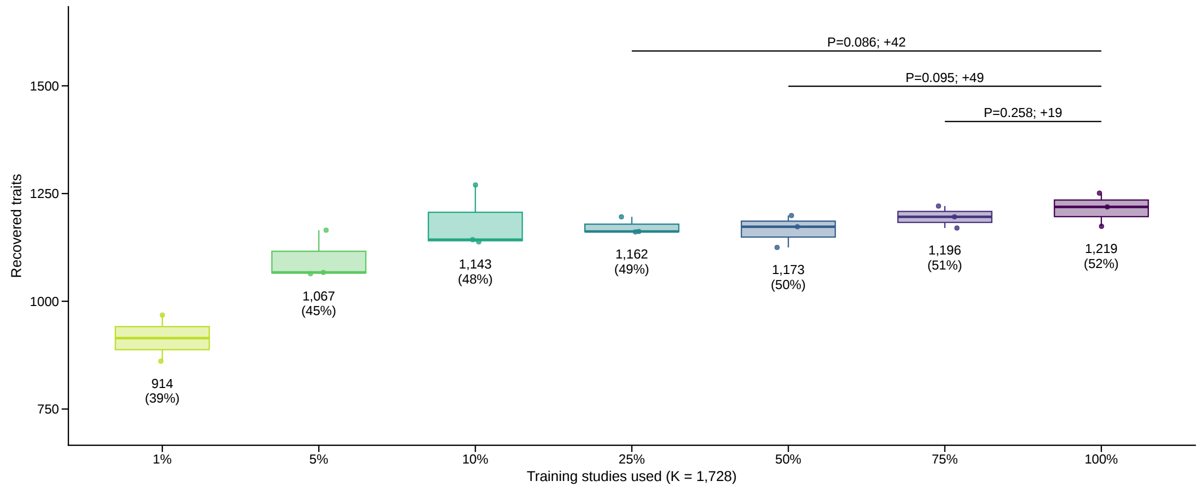


**Supplementary Figure 5: Trait recovery at a fixed number of gene modules across training sizes.** Traits recovered at FDR <0.05 by unsupervised CLAMPbase models retrained on each ARCHS4 subsample at $K$ = 1,728 ($n$ = 3 subsamples per fraction, except $n$ = 2 at 1%; Methods) and by the three CLAMPbase models trained on the complete ARCHS4 compendium (100%), whose estimated number of modules is 1,728, labelled as counts and as a percentage of 2,366 tested traits. Colours identify training fractions, as in Figure 3; boxes, median and interquartile range; whiskers, observations within 1.5 interquartile ranges; points, independent models; brackets, the mean gain of the 100% models over the labelled 75%, 50% and 25% fractions and one-sided Student's *t*-test *P* values. FDR, Benjamini-Hochberg false discovery rate.

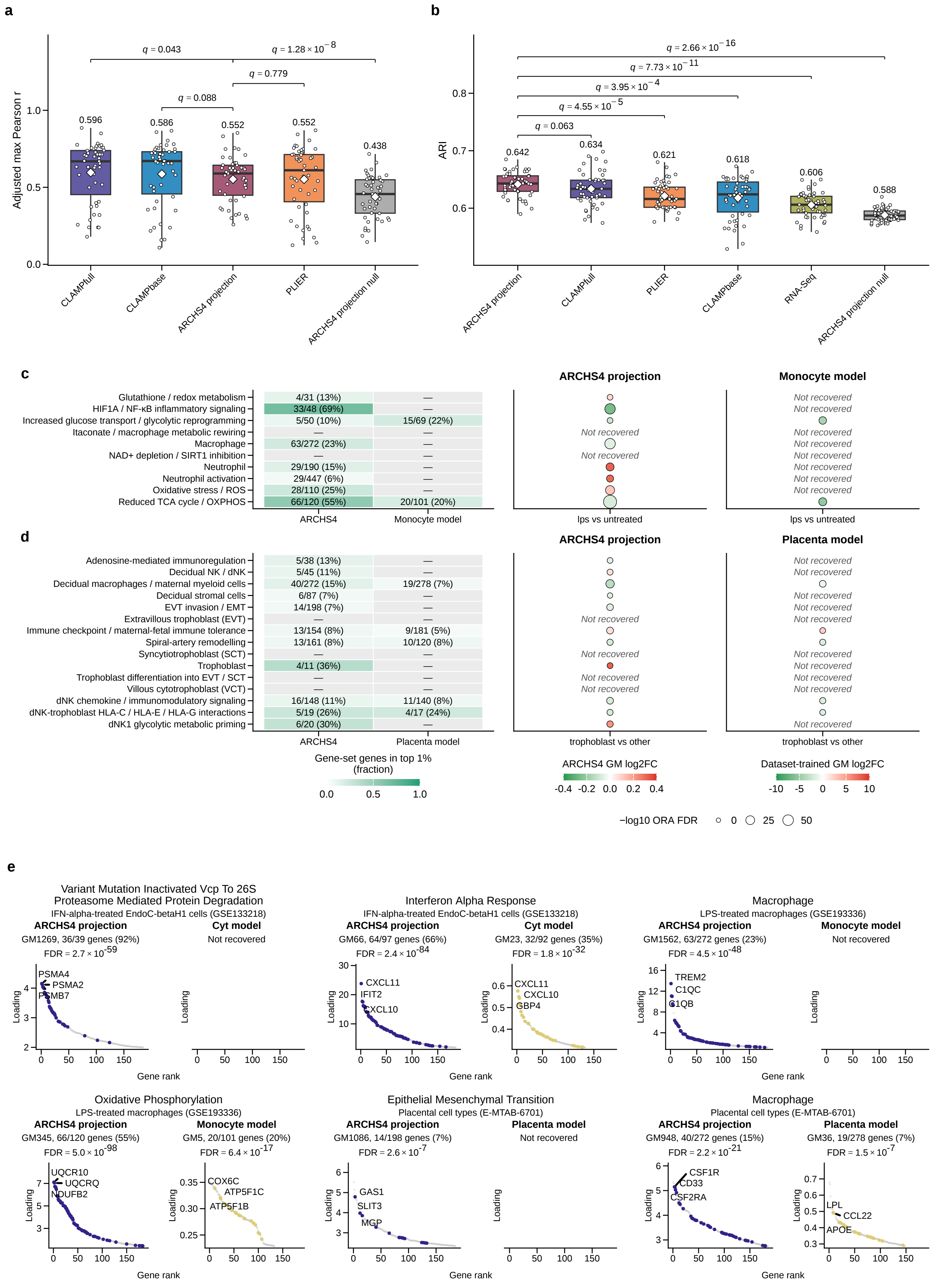


**Supplementary Figure 6: Projection benchmarks, recovery of pathways, processes and cell types, and gene loadings. a**, Cell-type proportion recovery in six pseudobulk cohorts ($n$ = 47 dataset-cell-type targets) by ARCHS4 projection and by dataset-trained methods fitted on each cohort, measured as the adjusted maximum Pearson correlation: the maximum across all 1,728 GMs for projection, or under greedy one-to-one matching for dataset-trained methods (Figure 2a), minus the median of 10,000 cell-proportion permutations. The null projection shuffles ARCHS4 loading rows across genes (mean of ten permutations). Points, targets; boxes, median and interquartile range; whiskers, 1.5 interquartile ranges; diamonds and numbers, means; brackets, paired two-sided Wilcoxon signed-rank tests, Benjamini-Hochberg-corrected ($q$). **b**, GTEx subtissue recovery measured by adjusted Rand index for projection into 578 random ARCHS4 GMs (the number of GMs in the dataset-trained GTEx models; a different subset in each repetition), dataset-trained methods, the RNA-Seq baseline and the null projection of all 1,728 GMs ($n$ = 50 clustering repetitions). Points, repetitions; boxes, whiskers, diamonds and numbers as in **a**; brackets, unpaired two-sided Wilcoxon rank-sum tests, Benjamini-Hochberg-corrected ($q$). **c**,**d**, Recovery of pathways, processes and cell types by ARCHS4 projection and by a CLAMPfull model fitted on the same dataset (Monocyte or Placenta model) in LPS-treated macrophages (**c**) and first-trimester placental cell types (**d**). Left, gene-set members among the top 1% of loadings of the matched module over members in the model's gene universe (percentage); fill, that fraction; em dash, not recovered. Right, module log2 fold change for the indicated contrast (fill; red, higher; green, lower; scale specific to each model type) and -log10(ORA FDR) (size); *Not recovered*, no module enriched for a matched gene set at ORA FDR <0.05. Targets, gene sets and supporting evidence are listed in Supplementary Table 2. **e**, Two examples per dataset, including the interferon-alpha response of Figure 4b, comparing ARCHS4 projection with the CLAMPfull model fitted on each dataset (Cyt, Monocyte or Placenta model): members of the matched gene set ranked by loading in the ARCHS4 (purple) and dataset-trained (ochre) module, other genes in grey, top three members labelled. Titles give the gene set and dataset; above each plot, the module, gene-set members among its top 1% over those in the gene universe (percentage) and the ORA FDR. LPS, lipopolysaccharide; EVT, extravillous trophoblast; SCT, syncytiotrophoblast; VCT, villous cytotrophoblast; dNK, decidual natural killer cell; GM, gene module; ORA, over-representation analysis; FDR, Benjamini-Hochberg false discovery rate.

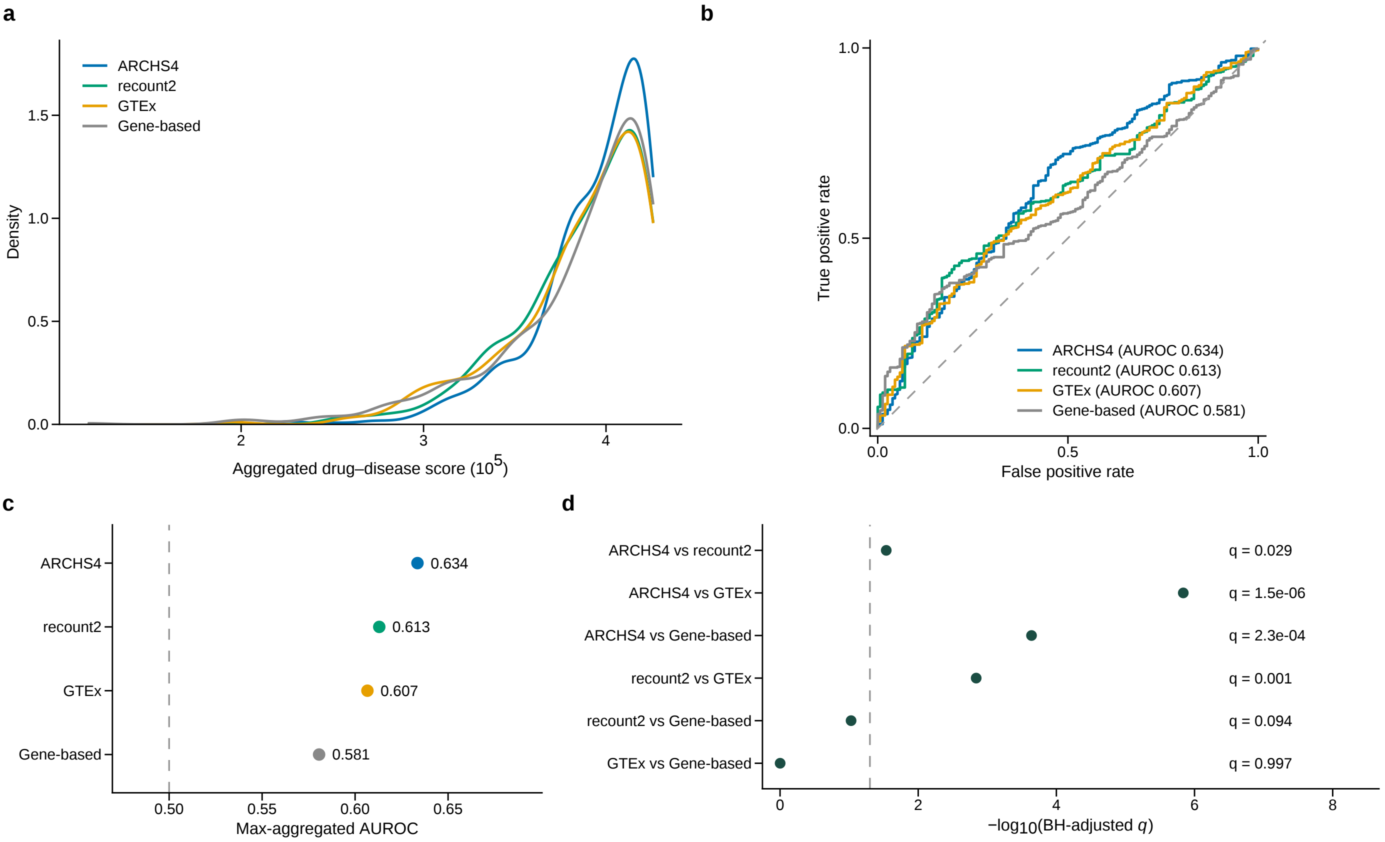


**Supplementary Figure 7: Drug-disease prediction and all pairwise method comparisons. a**, Kernel density of the aggregated drug-disease score for 685 labelled drug-disease pairs from a manually curated gold standard (531 pairs in which the drug is used to treat the disease and 154 in which it is not) spanning 337 drugs and 57 diseases, scored in the ARCHS4, recount2 and GTEx CLAMP models and with a gene-based approach applying the same scoring rule to individual gene associations; ranks were averaged across the top 5, 10, 25 and 50 modules (or 50, 100, 250 and 500 genes) and maximized over the 49 GTEx tissues (Methods); colours identify the four methods. The same pairs, methods and colours are used in **b** and **c**. **b**, Receiver operating characteristic curves for the scores in **a**; the legend gives each method's AUROC and the dashed line marks random expectation. **c**, AUROC of the scores in **a** for each method (values beside the points); dashed line, random expectation (AUROC = 0.5). **d**, BH-adjusted $q$ values ($-\log_{10}$ scale; exact values on the right) for all six pairwise comparisons of AUROC computed within each tissue for the fixed-threshold scores of Figure 4c ($n$ = 49 tissues), from paired one-sided Wilcoxon signed-rank tests with BH correction across the six comparisons; dashed line, $q$ = 0.05. AUROC, area under the receiver operating characteristic curve; BH, Benjamini-Hochberg.

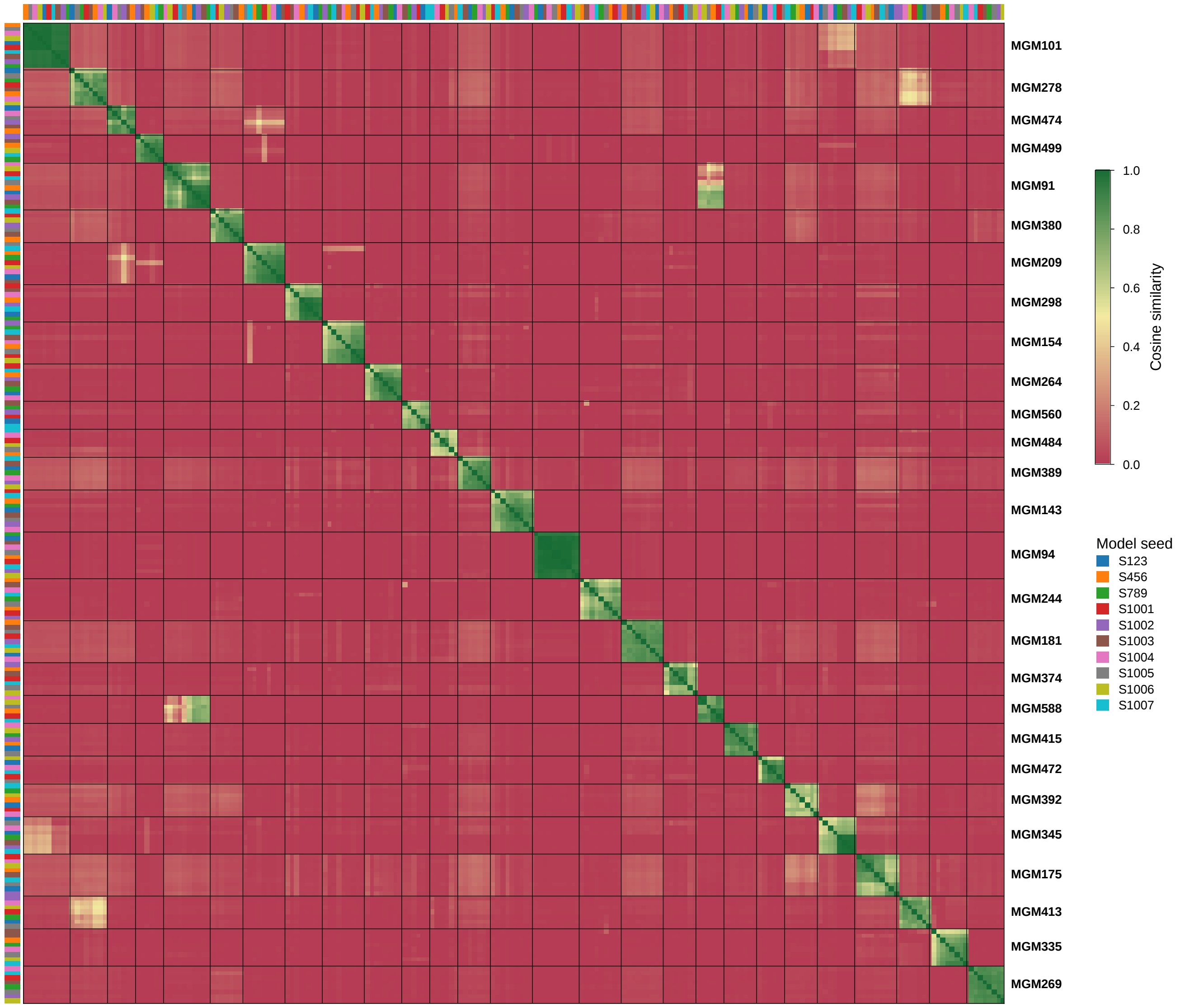


**Supplementary Figure 8: Lipid-screen MGMs reproduced across ten ARCHS4 models.** Cosine similarity between the gene-loading vectors of the member modules of the 27 MGMs enriched for a high-confidence HepG2 lipid-screen gene set (fgsea adjusted *P* <0.05, minimum across ten repetitions). MGMs were obtained from the 1,728 modules of each of ten CLAMPfull models trained on ARCHS4, the model used in the other analyses and nine replicates with different random initializations (colour strips, model seed), after removing modules flagged by the technical-noise screen (Supplementary Table 3), by complete-linkage agglomerative clustering on cosine distance cut at a similarity of 0.5 and retaining clusters with modules from at least six of the ten models (618 MGMs). Blocks, MGMs ordered by adjusted *P*; modules within a block ordered by complete-linkage clustering. Of the 27 MGMs, 19 were associated with a trait at FDR <0.05, 13 were also enriched for a canonical pathway, and the four with a lipid-related pathway are shown in Figure 5 (MGM91, MGM175, MGM335 and MGM588); all statistics and selection flags are listed in Supplementary Table 4. The enrichment threshold uses the minimum adjusted *P* value across repetitions and does not control the false discovery rate across the full screen, so the MGMs are prioritized candidates rather than confirmatory findings. MGM, meta-gene module; FDR, Benjamini-Hochberg false discovery rate.

### Supplementary tables

**Supplementary Table 1. Traits evaluated for genetic-trait recovery.** The 2,366 PhenomeXcan traits tested in the training-size, fixed-rank and complete-compendium analyses (Figure 3 and Supplementary Figure 5), with trait type, source and GWAS sample sizes (total, cases and controls); per-model trait recovery counts are provided as Figure 3 source data.

**Supplementary Table 2. Curated targets for the external-dataset projection comparison.** The 23 pathways or cellular processes and 10 cell types curated from the three source studies (Figure 4b and Supplementary Figure 6c,d), with the non-transcriptomic measurements that each study reported for the target and, for the ARCHS4 projection and the dataset-trained CLAMPfull model, whether the target was recovered, the matched module and gene set, and the ORA FDR.

**Supplementary Table 3. Technical-noise screen of the ten ARCHS4 models.** Gene-level and sample-level technical-noise flags and the cross-validated variance of sample scores explained by technical covariates for all 17,280 modules of the ten models (1,324 flagged; 116 of the 1,728 modules of the canonical reference), with per-model counts.

**Supplementary Table 4. Lipid-screen MGMs across ten ARCHS4 models and their selection.** The 618 MGMs reproduced in at least six of ten CLAMPfull canonical ARCHS4 models, with their member modules, the minimum and mean cosine similarity between members, fgsea results for the lipid-increasing and lipid-decreasing screen sets (minimum adjusted *P* across ten repetitions, NES and the CRISPR-Cas9 screen genes driving the enrichment), all screen genes among the top 1% of loadings and the cumulative selection flags (screen enrichment, trait association, canonical pathway, screen gene among the top 1% of loadings undetected by single-gene TWAS, lipid-related pathway); a second sheet gives the number of MGMs passing each step and a third summarizes the four MGMs of Figure 5 (models, screen genes, top genes, pathways, CellMarker and Azimuth cell types, ARCHS4 sample context, traits and genes with S-MultiXcan FDR <0.05).

**Supplementary Table 5. PhenoPLIER trait associations of the Figure 5 MGMs.** GLS associations of the four MGMs with the 2,366 PhenomeXcan traits, with trait sample sizes (samples, cases and controls), FDR and the disease to which each trait shown in Figure 5 was assigned.

**Supplementary Table 6. Over-representation analysis for the Figure 5 MGMs.** All terms enriched (ORA FDR <0.05) among the top 1% of genes of the four MGMs in GO Biological Process, canonical pathways, CellMarker, Azimuth and ARCHS4 cell lines, with overlap statistics, overlapping genes and the pathways shown in Figure 5.

**Supplementary Table 7. LINCS L1000 drugs scored on the Figure 5 MGMs.** For each of the four MGMs, all 1,170 LINCS L1000 drugs ranked by their standardized score on the MGM's top 1% of genes, with the mean z-score, permutation z-score, permutation *P* value and FDR, and the drug shown in Figure 5.

**Supplementary Table 8. Single-gene S-MultiXcan associations of the Figure 5 MGM genes.** S-MultiXcan z-scores, *P* values and FDR of the top 1% of genes and the screen genes of each MGM for the traits shown in Figure 5, with each gene's loading rank.
